# A Quantitative Two-Channel Genetic Reporter for Selenocysteine Biosynthesis and Incorporation

**DOI:** 10.64898/2026.08.09.743795

**Authors:** Andrew Gilmour, Qiyao Wei, Jessica Hellinger, Devon L. Kulhanek, Zach Jansen, Katelyn M. Baumer, Jennifer S. Brodbelt, Ross Thyer

## Abstract

Selenocysteine (Sec), the 21^st^ amino acid, is a rare non-canonical amino acid that represents an attractive target for protein engineering due to its desirable chemical properties such as high affinity for metals, strong nucleophilicity, and reversible covalent bond formation. To bypass the natural constraints on Sec placement within proteins, several strategies have been developed to rewire the native translational machinery to enable site-specific incorporation. However, these usually abolish the quality control mechanism that excludes the serine-charged selenocysteinyl-tRNA (Ser-tRNA^Sec^), the immediate biosynthetic precursor, from translation resulting in heterogenous protein species. This challenge is confounded by a lack of genetic tools to accurately report the selenylation state of the tRNA pool as most are blind to competing process of Ser incorporation, which can only be observed using analytical methods. To resolve this issue, we have developed a new fluorescent reporter, <u>Se</u>lenocysteine <u>A</u>djusted <u>R</u>atiometric <u>Ch</u>romophore (SeARCh), which exhibits two distinct spectral outputs dependent on the incorporation of either Ser (red) or Sec (green). Using SeARCh, we define several factors which influence the observed Sec:Ser ratio and construct a new hybrid biosynthetic pathway with improved performance, achieving 90% Sec incorporation. Furthermore, SeARCh displays unusually complex mass spectra due to the isotope distribution of selenium and heterogenous nature of the protein in solution and we report specific methods to account for this behaviour and precisely quantify the rare Ser-containing species found at high Sec incorporation efficiencies. Our findings suggest that the equilibrium between selenoprotein and tRNA^Sec^ expression levels is a key driver of incorporation efficiency and implies a process that is broadly biosynthetically constrained. Collectively these tools represent a significant advance in the metrology of selenocysteine biosynthesis and incorporation and can be used to inform and standardize future engineering efforts.

## Main

Non-canonical amino acids (ncAAs) that provide access to new chemistries beyond the standard 20 proteinogenic amino acids are invaluable tools for interrogating and expanding the function of proteins. Selenocysteine is a rare, naturally occurring ncAA with an unusual mosaic distribution in Nature. While it is structurally similar to cysteine, its unique selenol side chain yields distinct chemical and biophysical properties including a significantly lower pKa (5.2 vs 8.3 for free Cys), enhanced nucleophilicity, and the ability to form diselenide bonds which possess a lower reduction potential than their disulfide analogues(1–6). However, while some selenoproteins can be purified from their native hosts, efforts to express recombinant or synthetic selenoproteins have been limited by the complexity of biosynthetic and translational machinery needed to support introduction of a 21^st^ amino acid. As all sense codons within the standard genetic code are fully assigned, Sec is introduced into proteins by recoding a UGA stop codon. This process competes with termination of protein synthesis mediated by the host peptide release factors and in bacteria requires a stem-loop RNA structure (the SECIS element) adjacent to the UGA codon, within the coding sequence, to direct stop codon reassignment (**Figure 1a**)(7). In addition, Sec is unique in lacking a dedicated aminoacyl-tRNA synthetase and does not exist as a free amino acid. Instead, it is synthesized in place on the selenocysteinyl-tRNA which is pre-charged with serine (Ser) by the host seryl-tRNA synthetase. These inefficiencies and the imposed sequence constraints render the native machinery impractical for the synthesis of new selenoproteins.

**Figure 1.**
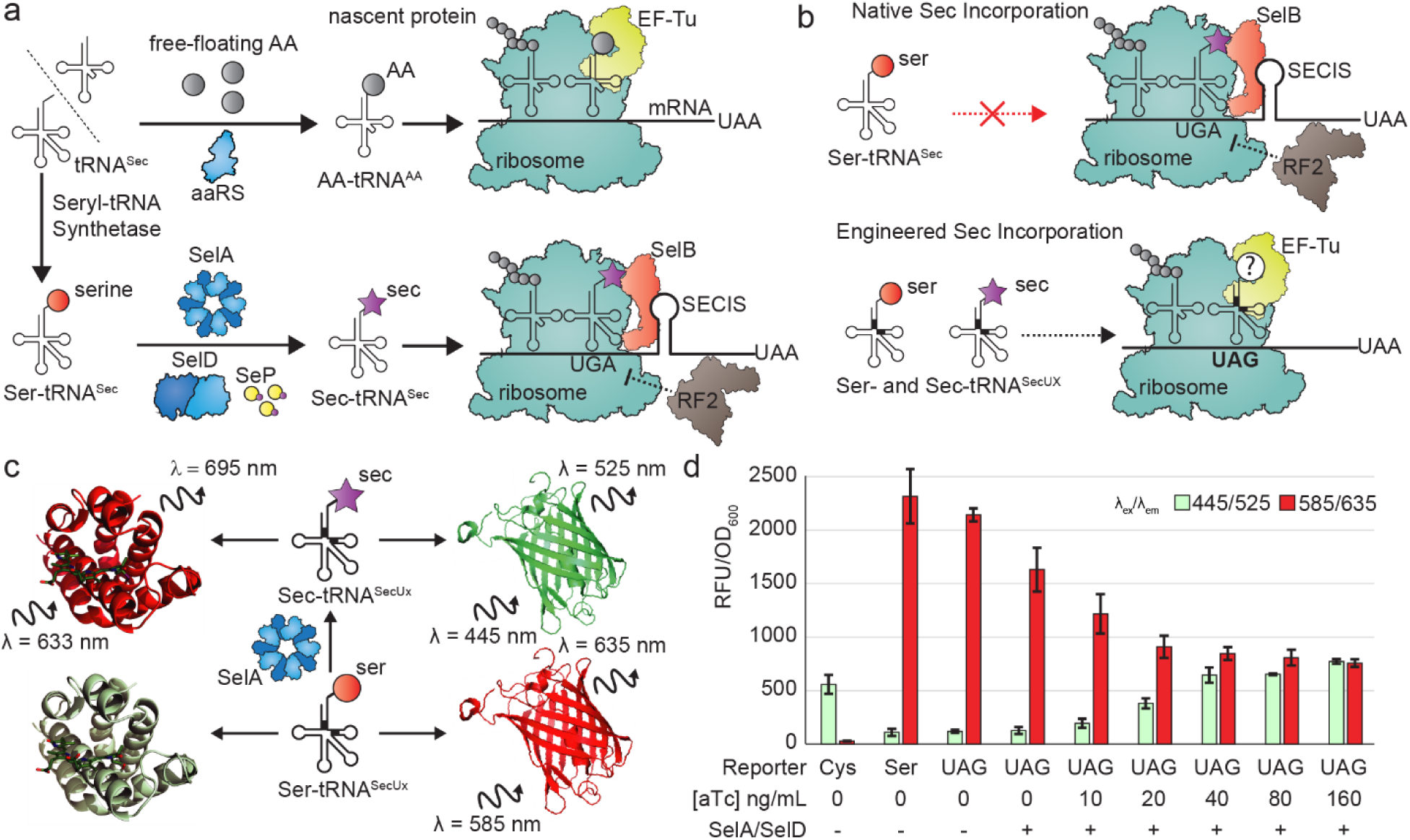
Selenocysteine biosynthesis and validation of the SeARCh reporter. (a) Schematic of canonical translation (top) and native bacterial Sec incorporation (bottom). tRNA^Sec^ must first be charged with serine before conversion to Sec-tRNA^Sec^ and then association with SelB and the SECIS element to recode a UGA stop codon. (b) Native bacterial Sec incorporation compared to engineered Sec incorporation. Ser-tRNA^Sec^ is not a substrate for SelB and thus will not participate in native translation. Engineered tRNA^SecUX^ (marked with block boxes) interacts with EF-Tu, eliminating the need for SelB/SECIS but also enables Ser-tRNA^SecUX^ to serve as a substrate for translation. (c) Sec-SmURFP (left) is a first-generation Sec-dependent genetic reporter that only yields a fluorescent signal when Sec is incorporated. SeARCh (right) represents a second-generation Sec-dependent reporter that generates two independent fluorescent phenotypes in response to Sec (green) or Ser (red) incorporation. (d) Validation of the SeARCh reporter using tRNA^SecUX^ presented as paired red and green fluorescence. Absence of SelA and SelD (pair 3) results in serine incorporation while increasing expression of the biosynthetic enzymes (SelA/D) via an aTc induction gradient (pairs 4-9) results in a dose-dependent increase in green and decreases in red fluorescence. Cys and Ser variants serve as theoretical incorporation maxima in their respective channels. (n=3, ±std).

Two recent advances have overcome these challenges and enabled researchers to readily generate novel selenoproteins. Firstly, genomically recoded *E. coli* strains with unassigned codons and lacking release factor 1 allow Sec to avoid competing with the termination machinery, greatly increasing the yield of recombinant selenoproteins(8, 9). Secondly, several new biosynthetic pathways have been developed which leverage engineered tRNA^Sec^ species which are substrates for the canonical translation elongation factor EF-Tu, rather than the selenocysteine-specific elongation factor SelB(10–13). The requirement for SelB to associate with the cis-acting RNA structure is responsible for the sequence constraints on Sec incorporation, while EF-Tu can support truly site-specific incorporation(7, 14, 15). However, while these advances have unlocked the ability to synthesize new selenoproteins, SelB provides a critical gate-keeping mechanism to exclude serylated tRNA^Sec^ from participating in translation which EF-Tu cannot replicate (**Figure 1b**). This results in spurious and highly variable incorporation of Ser, imposing a serious challenge for characterization as the Ser- and Sec-containing isoforms cannot be physically separated nor their relative contributions resolved during biochemical characterization.

Strategies to tackle this issue, include more efficient biosynthesis to shift the equilibrium of the global tRNA^Sec^ pool towards the selenylated species, or re-establishing translational quality assurance to exclude Ser through engineered SelB variants(16). However, applying high-throughput molecular methods to this problem is difficult, given the challenges associated with measuring Sec incorporation in living cells. In contrast to other ncAAs, which can be measured using genetic reporters which rely solely on stop codon suppression, this process cannot discriminate between Sec and Ser incorporation events. While a variety of selenocysteine-specific reporters have been developed which effectively read the side chain of the incorporated amino acid, including selectable markers(11), fluorescent proteins(3), and flexible intein-based systems(17), none quantify Ser incorporation. Thus, the overall ratio of Sec:Ser incorporation must be determined using analytical methods. Additionally, the diversity of these genetic reporters, target selenoproteins and their expression conditions, and mass spectrometry methods also make it difficult to directly compare literature reports and draw broad conclusions about factors which influence incorporation efficiency.

Here, we report the development of a new ratiometric fluorescent reporter protein, the <u>Se</u>lenocysteine <u>A</u>djusted <u>R</u>atiometric <u>Ch</u>romophore or SeARCh, which yields two distinct spectral outputs that correspond to either Ser (red) or selenocysteine (green) incorporation (**Figure 1c**). SeARCh provides an unparalleled ability to visualize the selenylation status of the tRNA^Sec^ pool in near real time, enabling optimization of the Sec biosynthetic machinery as well as fundamental investigation of both molecular and bioprocess factors which influence incorporation efficiency. Importantly, this protein displays unusual characteristics during mass spectrometry which necessitated the development of specific methods to resolve the discrete Sec-containing fragment while accurately accounting for relevant low-abundance SeARCh proteoforms. This revealed differences in measurement of Sec incorporation efficiency depending upon the data processing workflow and may represent another unintended source of variation. Using SeARCh, we have identified several parameters which strongly influence Sec incorporation efficiency, including the addition of a new biosynthetic step, establishing its utility as a precision tool for the metrology of selenocysteine.

## Results

### SeARCh: a novel, two-channel reporter for selenocysteine incorporation

To develop a dynamic reporter capable of quantifying incorporation of both Sec or Ser, we sought to identify a protein scaffold that exhibits distinct, measurable phenotypes when alternating between cysteine (Cys) and Ser at a single residue. We identified the red fluorescent protein, mKate, as a candidate due to two reported mutations, Ser143Cys and Ser158Ala, that yield a pH-dependent chromophore shift from red (*λ*_ex_: 585 nm, *λ*_em_: 635 nm) to green (*λ*_ex_: 445 nm, *λ*_em_: 525 nm)(18). This reversible color transition is based on changes in protonation state of its *cis*-chromophore and is pronounced at physiological pH(18). We replicated this result by constructing two versions of mKate2 A45V with either Cys143 or Ser143. These variants displayed markedly different signal intensities in their respective channels, which matched the previously reported differences in brightness (**Figure 1d**).

To investigate whether this reporter could provide a dual-channel response based on the changes in the ratio of Sec:Ser at residue 143, we assembled a minimal Sec biosynthetic and incorporation pathway using a modular genetic framework to allow precise control over the individual genetic elements. This comprised a series of modular transcriptional and translational control elements, including bi-cistronic design (BCD) ribosome binding sites (RBS) and insulating bidirectional synthetic terminators, which serve to minimize any contextual influence(19). To facilitate this assembly scheme, we first refactored the engineered selenocysteinyl-tRNA variant tRNA^SecUX^ to ensure compatibility with common Type IIS restriction enzymes(11). This necessitated removal of a native Esp3I restriction site within the D-stem. Based on alignments with orthologous tRNA^Sec^ sequences from *Yersinia enterolitica* and *Enterobacter sp.638*, we introduced new C_14_-G_22_ (*Yer*) and A_11_-T_25_ (*Ent*) base pairs (**Figure S1a-b**). We validated that this minor structural change did not compromise tRNA^SecUX^ function using Seleno-SmURFP, where incorporation of Sec yields red fluorescence by enabling ligation of the biliverdin chromophore (**Figure S1c**)(3). The C_14_-G_22_ base pair had no detectable effect of activity and activity and the new tRNA variant was designated tRNA^SecUXY^.

The genomically recoded *E. coli* strain B95.ΔA with all four dedicated Sec biosynthesis genes deleted (Δ*selABCD*) was used as the host for all experiments. We hypothesized that tight control over the expression of the Sec biosynthesis enzymes and the reporter would allow us to precisely measure whether changes in biosynthetic flux are linked to a higher green to red ratio, consistent with greater Sec incorporation. A two-plasmid system was developed consisting of a reporter plasmid encoding SeARCh (Cys/Ser/UAG143) under the control of the inducible P_BAD_/AraC regulon, and a Sec biosynthesis plasmid encoding *selA*, *selD*, and tRNA^SecUXY^, where the two biosynthetic enzymes were placed under the control of a tightly regulated P_LtetO_/TetR system. The tRNA^Sec^ was expressed from the strong constitutive *E. coli* tRNA promoter *leuQ*. Initial results showed that in the absence of SelA and SelD, tRNA^SecUXY^ was efficiently charged with Ser, leading to near-complete UAG codon suppression and a red fluorescent phenotype (**Figure 1d**). However, upon induction of the two biosynthetic enzymes, we observed a dose-dependent decrease in red fluorescence accompanied by a corresponding increase in green signal, confirming that: (i) Sec incorporation supports formation of the green chromophore, and (ii) that greater selenylation of the tRNA pool drives a ratiometric change in the spectral output. Successful detection of both Sec and Ser incorporation events led us to name this reporter (mKate2 A45V S143U) the <u>Se</u>lenocysteine <u>A</u>djusted <u>R</u>atiometric <u>Ch</u>romophore or SeARCh.

### Investigating the rate-limiting step in Sec biosynthesis

Given the inherent challenges monitoring the intracellular Sec:Ser ratio using traditional methods, systematic investigation of factors affecting Sec incorporation efficiency is lacking. While kinetic studies have confirmed the catalytic rate of SelA is slow, and far below that of an aminoacyl-tRNA synthetase, it is not clear whether it actually limits in the pathway in practice or whether excess SelA is deleterious due to unproductive tRNA sequestration(20–22).

We hypothesized that if SelA were the rate-limiting step, an induction gradient would yield a linear increase in Sec incorporation and thus green fluorescence when SelD and tRNA^SecUXY^ expression were held constant and in excess. Conversely, if SelA was not the limiting factor, green fluorescence would rapidly saturate. To test this theory, we expressed SelD and tRNA^SecUXY^ constitutively while placing *selA* under the strong P_LtetO_ promoter. An increase in green fluorescence and concomitant decrease in red fluorescence was observed across the full induction range (0–80 ng/mL anhydrotetracycline (aTc)), consistent with a limiting step (**Figure 2a**). However, the corresponding circuit titrating SelD expression (with SelA held constant) yielded a nearly identical result (**Figure 2b**). These data suggest that neither enzyme serves as a clear bottleneck under these conditions; rather, the pathway appears to benefit from the simultaneous upregulation of both components. This would be consistent with structural studies which postulate the formation of a functional SelA:tRNA^Sec^:SelD complex and implies the stoichiometry between SelA and SelD is in important consideration.

**Figure 2.**
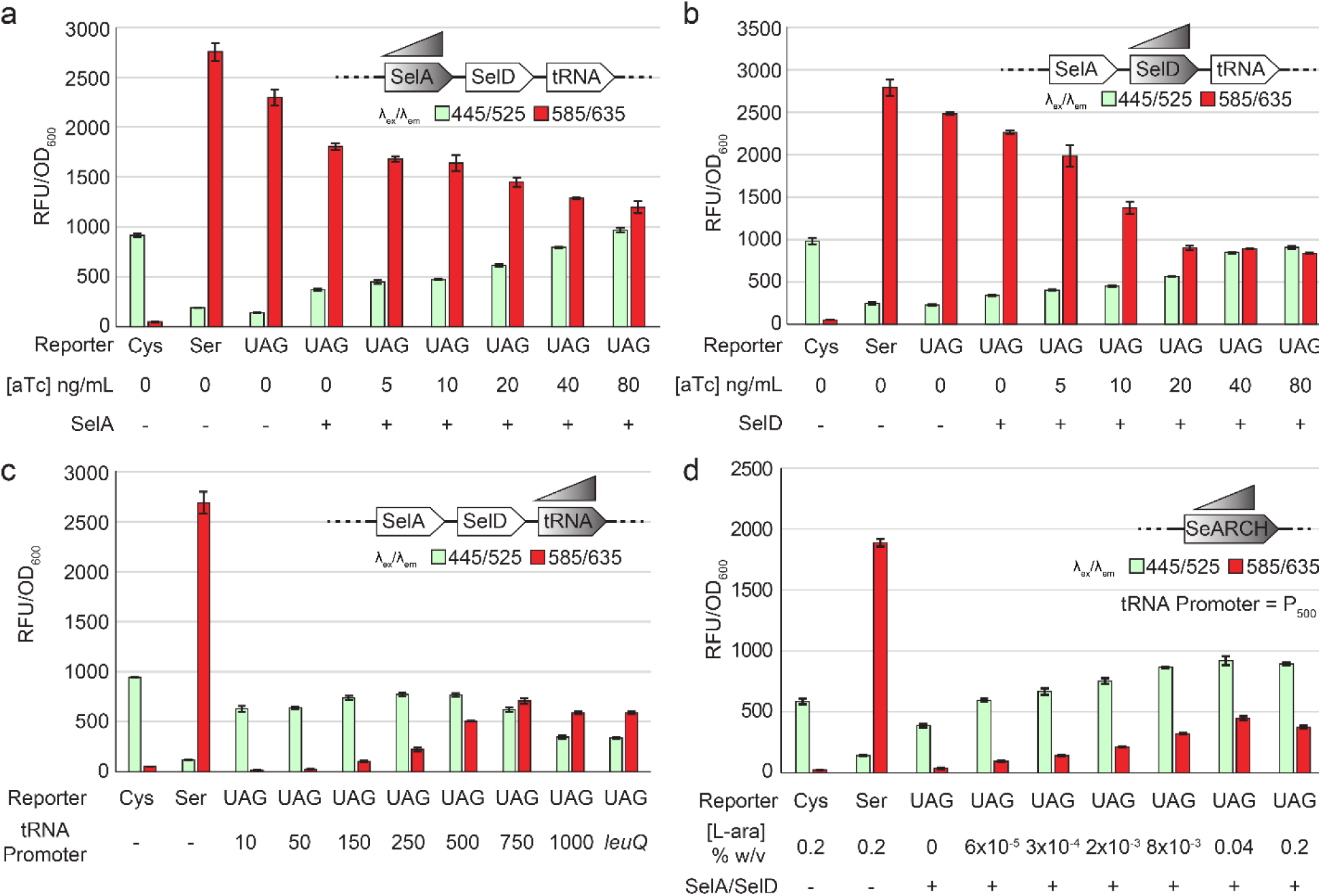
Titration of Sec biosynthesis machinery and SeARCh reporter. Two channel fluorescence from the SeARCH reporter where either SelA (a) or SelD (b) is titrated using a gradient of aTc (0-80 ng/mL) while the other enzyme is held constant and tRNA^SecUXY^ is expressed using a strong constitutive promoter. (c) Fluorescence assay where tRNA^SecUXY^ is expressed using a gradient of constitutive promoter strengths (10-1000, arbitrary units) and the *E. coli leuQ* tRNA promoter. Stronger promoters result in greater red fluorescence suggesting that excess tRNA^SecUXY^ is driving higher rates of serine incorporation. (d) Titration of the SeARCh reporter using L-arabinose while the Sec biosynthetic machinery is held constant. Low induction of the reporter yields high Green:Red ratios (∼5:1) while increasing induction strength lowers the ratio to (∼2:1). All panels (n=3, ±std).

While tRNA^Sec^ has been the focus of engineering efforts aimed at reducing the complexity of the Sec biosynthetic and incorporation pathway, how changes in tRNA expression level impact incorporation fidelity is unknown. However, kinetic evidence indicates that the aminoacylation of tRNA^Sec^ with Ser by the host SerRS occurs significantly faster than the subsequent selenylation step to form Sec-tRNA^Sec^. Excessive tRNA^Sec^ expression may promote Ser misincorporation through the formation of a large pool of serylated tRNA which cannot be completely converted by the slower terminal biosynthetic step. To investigate the relationship between tRNA expression and Sec incorporation fidelity, we used a library of constitutive promoters spanning a 100-fold range (10 to 1,000 relative units) (**Figure 2c**). Our results reveal that tRNA^SecUXY^ expression has a profound impact on fidelity with high-level constitutive expression being a key driver of Ser misincorporation. Notably, the original *leuQ* promoter performed similarly to our strongest test variants (P_1000_), suggesting that Sec biosynthetic pathway designs that favored high levels of tRNA expression to compete with termination before the widespread use of genomically recoded termination-deficient strains are biased towards to high Ser misincorporation.

### Selenoprotein expression affects Sec incorporation fidelity

These initial experiments imply that the terminal step of Sec biosynthesis is ultimately the limiting factor to achieving high Sec:Ser incorporation ratios. Based on this information, we would also expect that higher expression of a selenoprotein could deplete the Sec-tRNA^Sec^ pool at a faster rate than the biosynthetic machinery can replenish it, resulting in the more rapidly generated Ser-tRNA^Sec^ species entering translation. To test this hypothesis, we titrated expression of our SeARCh reporter while holding the expression of SelA, SelD, and tRNA^SecUXY^ constant. We paired the moderately strong P_500_ constitutive promoter with tRNA^SecUXY^ since it resulted in an intermediate green/red ratio at full induction of SelA/D, making it more sensitive to perturbations in the spectral output (**Figure 2d**). In agreement with this hypothesis, low-level expression of the SeARCh reporter results in high green-to-red ratio (∼5:1) while progressively stronger induction shifts the ratio towards red (∼2:1). This confirms that increased demand on the tRNA^Sec^ pool compromises translational fidelity and favours Ser misincorporation. We similarly evaluated SeARCh expression using a standard pET expression plasmid. Expression from the strong bacteriophage T7 promoter was anticipated to place further strain on the Sec biosynthetic pathway and we expected a significant shift towards predominantly Ser incorporation. A subset of the constitutive promoters was used to express tRNASec^UXY^ in conjunction with full induction of SelA and SelD. Our results show a dramatic increase in the red channel for the promoter strengths at P_150_ and above (**Figure S2**). These indicate that restricting either tRNA^Sec^ or selenoprotein expression can be effective strategies to improve the fidelity of Sec incorporation. In addition, we believe these phenomena, explained by differences in host strain (e.g. non-recoded, C321.ΔA, B95.ΔA), protein targets (e.g. hGPX-1, DHFR, HGH), and methods of expression (e.g. promoters, induction levels, plasmid copy number), account for the high variability in Sec:Ser incorporation ratios reported in the literature(3, 8–10, 13, 16).

### SeARCh aligns with traditional biochemical reporters

Beyond the dedicated biosynthetic and incorporation machinery encoded by the *selABCD* genes in bacteria and the requirement for serylation of tRNA^Sec^ by the host SerRS, the broader biosynthesis pathway remains to be determined. A critical gap in our understanding is how Sec biosynthesis interfaces with inorganic selenium metabolism, in particular, the identity of the Se species that serves as the substrate for SelD *in vivo* remains unknown. In previous work, the *E. coli* thioredoxin system, comprising thioredoxin (TrxA) and its partner reductase (TrxB), were proposed to catalyze the essential reductive assimilation of Se and it was demonstrated that disruption of either gene abolishes activity of the endogenous *E. coli* selenoprotein formate dehydrogenase H (FDH_H_)(23). To investigate whether the thioredoxin system is required for Sec biosynthesis in *E. coli*, we recapitulated the disruption of *trxA* and *trxB* and used the resulting strains to express the SeARCh reporter. Genes were disrupted using a CRISPR-associated transposon (CAST) system with two independent guides for *trxA* and *trxB* (**Figure S3**) (24, 25). Unexpectedly, we observed no difference in the spectral output of SeARCh when either *trxA* or *trxB* was deleted (**Figure 3a**). A direct assay of formate dehydrogenase activity using the redox-active dye benzyl viologen confirmed this result, suggesting that neither gene is essential for Sec biosynthesis (**Figure 3b**). We speculate that prior results are due to strain-specific factors and the fact that the thioredoxin system is integral to the cell’s redox biology more generally, including the assembly of iron-sulfur clusters. These observations highlight the value of SeARCh as a reporter for Sec biosynthesis and incorporation as the protein is fully decoupled from bacterial metabolism, requiring no co-factors or substrates, and folds and matures independently of host proteins.

**Figure 3.**
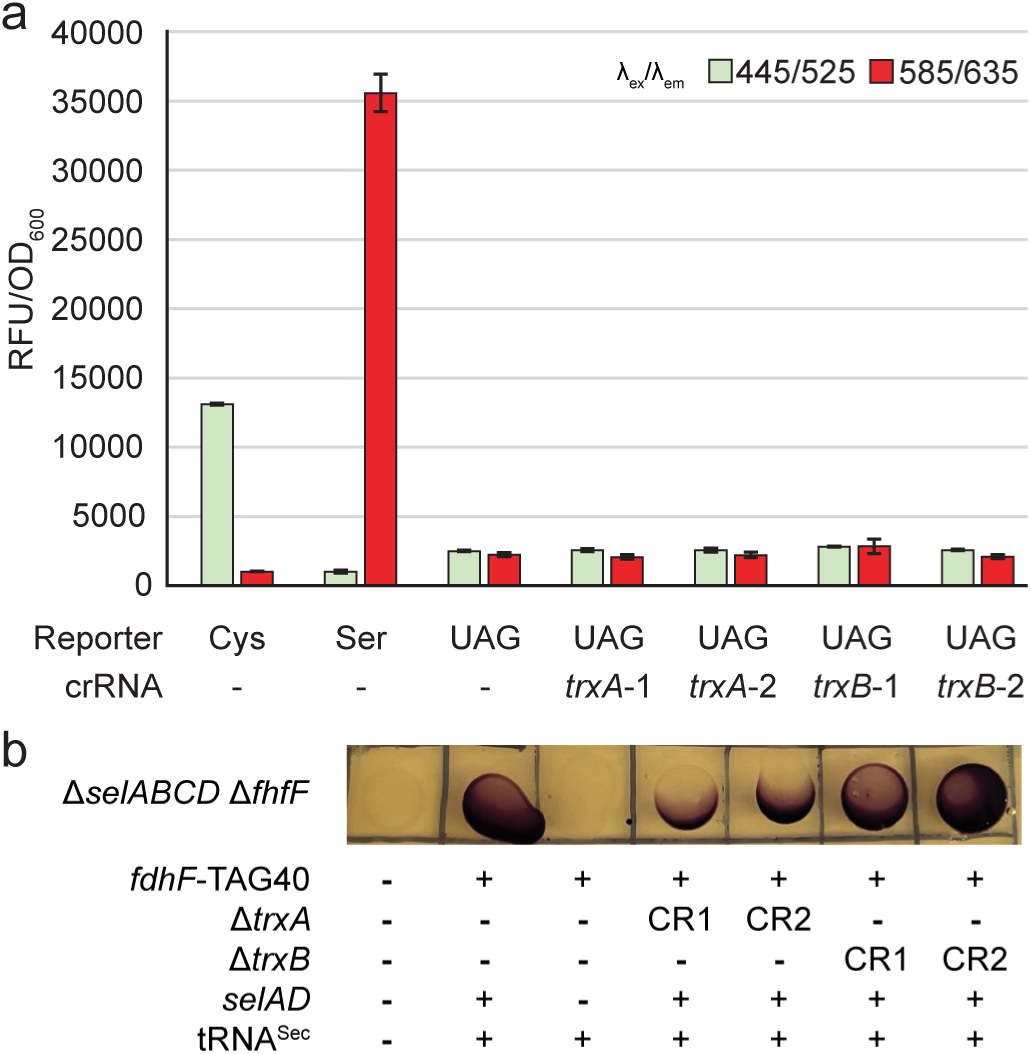
Evaluation of genetic disruption of *trxA* and *trxB* using SeARCh reporter. (a) Fluorescence assay evaluating SeARCh reporter fluorescence in conjunction with two independent genetic disruptions of *trxA* and *trxB* (denoted -1 or -2). Genetic disruptions result in equivalent ratios of Green:Red fluorescence compared to wild-type. (n=3, ±std). (b) Reduction of benzyl viologen by *E. coli* formate dehydrogenase H (*fdhF*) with a TAG stop codon at residue 40 is strictly dependent on Sec incorporation (representative colonies). No color indicates a lack of activity due to either the absence of FDH (cells only) or selenocysteine biosynthetic enzymes (tRNA^Sec^ + *fdfF*-TAG40). Disruptions of *trxA* or *trxB* did not abolish Sec incorporation.

Given this result, we investigated whether other genes involved in bacterial selenium metabolism would influence Sec biosynthesis and result in a loss or reduction of green fluorescence from the SeARCh reporter when inactivated. We disrupted an additional nine non-essential genes linked to sulfate transport (*cysP*), inorganic phosphate transport (*pitA*/*pitB*/*pstC*), cysteine desulfurases (*csdA*/*sufS*), and the glutathione system, including glutathione biosynthesis (*gshA*/*gshB*), and glutathione reductase (*gorA*), each using two independent guides (**Figure S4a-b**)(26–30). Our findings indicate that none of the genes evaluated are essential for Sec biosynthesis by themselves, and that early steps of bacterial Se metabolism are carried out by yet unidentified enzymes or there is widespread redundancy that obfuscates the contributions of individual genes.

### Bioprocess optimization of selenoprotein expression

The flexibility of the SeARCh reporter enables the interrogation of not only genetic architecture but also bioprocess conditions such as media composition, assay duration, and supplementation of inorganic selenium donors. We initially used only M9 minimal medium supplemented with 2.5 g/L yeast extract (M9YE) and 0.5% glycerol but observed significant growth defects under high-induction conditions. We hypothesized that a richer medium might yield greater biomass and counteract toxicity of highly active Sec biosynthesis. While media richness correlated positively with final OD_600_, the fluorescent outcomes revealed a complex relationship between growth and fidelity. Terrific Broth (TB) significantly increased biomass but provided very little increase in normalized fluorescence (**Figure S5**). At six hours, spectral differences across media were negligible; however, by 16 hours, distinct phenotypes emerged. M9YE yielded the lowest red fluorescence, suggesting superior Sec-incorporation fidelity, but suffered from the lowest overall protein yield. This observation further supports the connection between lower target protein expression and higher Sec incorporation fidelity. Interestingly, half strength TB (½TB) outperformed TB at 16 hours, exhibiting higher signal in both channels.

An often-overlooked parameter in tuning selenocysteine biosynthesis is the concentration of the inorganic selenium donor, typically Na_2_SeO_3_ (sodium selenite). To define the optimal supplementation regime, we evaluated a selenite gradient from 0 to 200 μM (**Figure S6**). In the absence of exogenous selenite, the fluorescence profile closely mirrored the Ser-only control, though a marginal increase in green signal suggested trace selenium levels in the rich media support basal Sec biosynthesis. We observed a dose-dependent increase in green fluorescence that peaked at 25 μM [Se]; concentrations exceeding this threshold led to a steep decline in both green fluorescence and optical density OD_600_. To pinpoint the transition between maximal productivity and metabolic stress, we performed a finer titration between 25 and 50 μM. This revealed that 30 μM Na_2_SeO_3_ provides an optimal balance under these media conditions, maintaining peak green fluorescence, while suppressing the red channel and achieving reasonable biomass. Consequently, we selected ½TB and 30 μM Na_2_SeO_3_ in combination with P_250_-tRNA^SecUXY^ and inducible SelA/D as our standardized expression conditions for further optimizations.

### An additional biosynthetic step can reduce serine misincorporation

While bacteria directly selenylate Ser-tRNA^Sec^ to yield the mature Sec-tRNA^Sec^ through the actions of SelA, in Archaea and Eukaryota this process proceeds via a phosphorylated intermediate (Sep-tRNA^Sec^). This is generated by a dedicated kinase, O-phosphoseryl-tRNA kinase (PSTK), and is strictly essential as SepSecS, which is functionally analogous to bacterial SelA, has no activity on Ser-tRNA^Sec^(31, 32). Addition of PSTK to engineered Sec biosynthesis pathways in *E. coli* has been investigated previously, under the assumption that Sep-tRNA^Sec^, which is a poor substrate for translation due to weak interactions with EF-Tu, is still a substrate for SelA. However, these experiments have not shown conclusively beneficial results. We hypothesized that a PSTK isoform from an organism that shares sequence motifs in its tRNA^Sec^ with our engineered tRNA^SecUXY^ might yield greater improvement (**Figure 4a**)(10, 33). While eukaryotic orthologs of PSTK primarily use the length and secondary structure of the D-stem to discriminate tRNA^Sec^ from tRNA^Ser^, archaeal PSTK orthologs have been found to recognize key acceptor stem base pairs (G_2_-C_71_, C_3_-G_70_) and minor identity elements in the D-arm and T-loop(32, 34).

**Figure 4.**
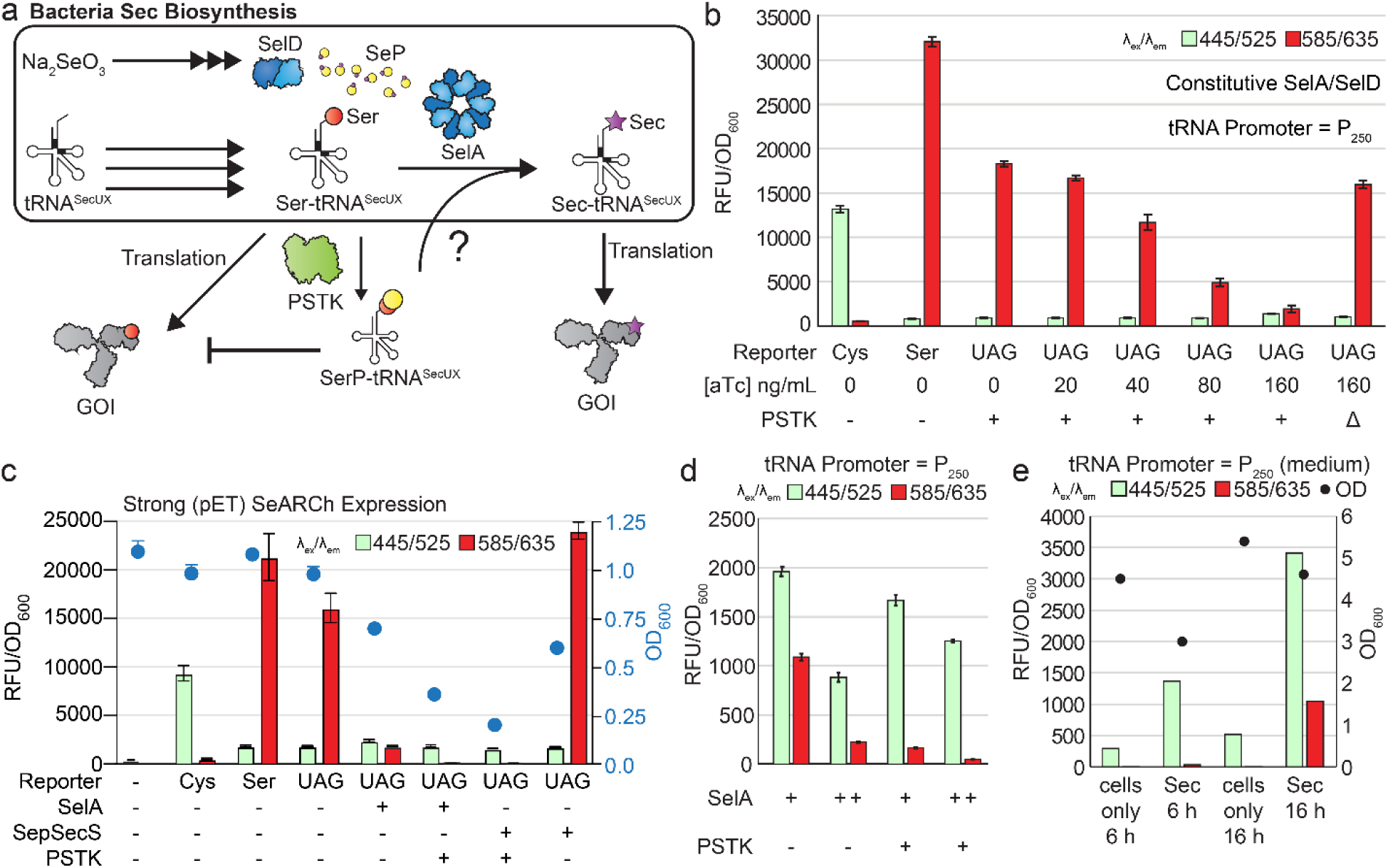
Expression of an archaeal PSTK can reduce serine incorporation. (a) Schematic of bacterial Sec biosynthesis pathway depicting potential interactions with PSTK and implications for Ser and Sec incorporation. (b) Fluorescence assay using pET-SeARCh reporter titrating expression of *Mk*PSTK via an aTc gradient. Expression of SelA/SelD/tRNA^SecUXY^ was biased towards serine incorporation and held constant. (c) Confirmation of activity of *Mk*SepSecS using pET-SeARCh reporter. Lack of activity in the absence of *Mk*PSTK indicates that *Mk*SepSecS activity is dependent on the generation of SerP-tRNA^SecUXY^ and that *Mk*PSTK is functionally active. Expression of both *Mk*PSTK and *Mk*SepSecS result in reduction of cell biomass as measured by OD600. (d) Fluorescence assay of SeARCh reporter with increased SelA by tuning BCD RBS strength (denoted by + or ++) improved the Green:Red ratio by ∼6x. SelA, SelD, and *Mk*PSTK are all under the inducible control of the PLteto/TetR while the tRNA^SecUXY^ is expressed constitutively with P250. (n=3, ±std). (e) Fluorescence assay of SeARCh reporter from 200 mL baffled flasks. 6-hour expression leads to lower OD600 and overall protein expression but results in a higher Green:Red ratio compared to 16 hours expression. (n=1, representative).

The PSTK from *Methanopyrus kandleri* (*Mk*PSTK) was identified as a potential candidate as several of the tRNA bases which are predicted to form contacts with the protein are maintained in *E. coli* tRNA^Sec^ and tRNA^SecUXY^. The *M. kandleri* enzyme has no close homologs, so we also evaluated two additional Archaeal PSTK, the *Methanocaldococcus jannaschii* (*Mj*PSTK), which has been investigated previously as a tool for improving Sec biosynthesis, and *Methanococcus aeolicus* (*Ma*PSTK) which has low sequence conservation with the other two candidates. We expressed these PSTK variants along with the core Sec biosynthetic pathway which we configured to favour high Ser incorporation to make potential changes in SeARCh spectral output more easily observed. Co-expression of the *Mk*PSTK showed a dramatic reduction in red signal while expression of *Mj*PSTK and *Ma*PSTK showed no change (**Figure S7**). To confirm this effect was solely due to *Mk*PSTK activity and not changes in the context of the Sec biosynthesis plasmid, we also constructed an inactive version by truncating the protein after residue L90 and found it performed no differently than a ΔPSTK control. To further establish that PSTK activity was responsible for the loss of red fluorescence, we titrated expression of *Mk*PSTK using a gradient of aTc and found that higher expression of *Mk*PSTK strongly reduced red fluorescence while slightly increasing signal in the green channel (**Figure 4b**). To confirm that PSTK was generating SerP-tRNA^Sec^ and not influencing Sec biosynthesis via another mechanism, we tested whether *Mk*SepSecS could complement deletion of *Ec*SelA (**Figure 4c**). Our results indicate that *Mk*SepSecS is functional in *E. coli*, but critically, only when *Mk*PSTK is also expressed. While PSTK expression results in very low red fluorescence, indicating strong suppression of Ser misincorporation, our initial expression context was accompanied by a significant decrease in biomass, indicating toxicity when this enzyme is highly expressed. Thus, expression of SelA, SelD and *Mk*PSTK was rebalanced to yield more robust cell growth and the highest possible Green:Red fluorescence ratio from the SeARCh reporter. Increasing the translation rate of SelA via a Bicistronic Design Ribosome Binding Site (BCD) improved the Green:Red ratio but was generally associated with a penalty to biomass and protein yield, with less fluorescence observed across both channels. Optimal PSTK expression was achieved using the weakest BCD available. In conjunction with a shorter 6 h induction of the reporter, this pathway configuration resulted in a Green:Red ratio exceeding (**Figure 4d-e**).

### Mass spectrometry of SeARCh

To experimentally validate that changes in fluorescence from the SeARCh reporter correspond to meaningful differences in Sec incorporation fidelity, we purified the SeARCh reporter expressed using a series of conditions empirically determined to yield different ratios of Green:Red fluorescence. A control condition in which the Sec biosynthesis enzymes were not induced was also included where we expected to observe only Ser incorporation. To ensure that the results were not unique to the SeARCh reporter and generalizable to other proteins, identical conditions were used to express sfGFP encoding a single UAG stop codon at a permissive position (S205UAG). SeARCh samples showed excellent correlation between increasing Green:Red ratio (**Figure 5a**) and selenocysteine incorporation based on intact mass analysis (**Figure S8**). A sample expressed using the strongest induction of the Sec biosynthetic enzymes (160 ng/mL aTc) yielded 91 mg.L^-1^ of protein with a ∼42/58 ratio of Green:Red fluorescence corresponding to 77% selenocysteine incorporation based on the relative abundances of the corresponding intact protein ions and the C-terminal fragment ions containing residue 81-249. This is consistent with the brighter fluorescence of the red chromophore and provides enhanced sensitivity at low levels of Ser incorporation where it is most important. The sfGFP S205UAG control samples showed near identical incorporation ratios, suggesting that optimizations made with the SeARCh reporter can translate to other proteins (**Figure S9**).

**Figure 5.**
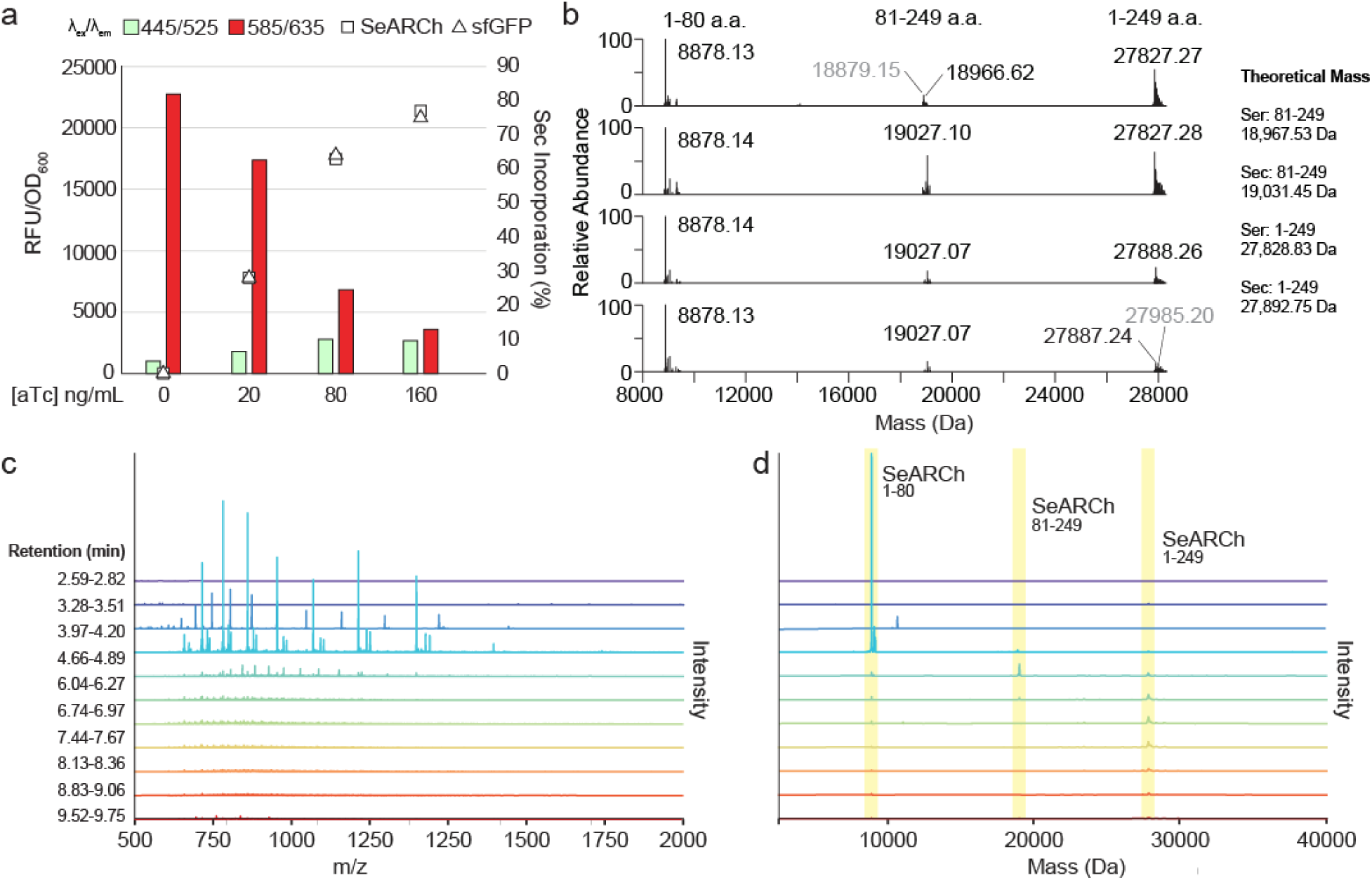
Mass spectrometry validation and optimization of SeARCh reporter. (a) Paired fluorescence and % Sec incorporation calculated based on MS1 analysis from cells expressing SeARCh using a strong T7 promoter. sfGFP S205UAG was expressed, purified, and analyzed under identical conditions and resulted in near identical incorporation efficiencies, confirming that conditions for Sec incorporation are transferable to different protein scaffolds. (n=1, representative) (b) Deconvoluted MS1 spectra obtained for 10 μM SeARCh (27.9 kDa) in 50:50 methanol:water with 0.1% formic acid. From top to bottom, SeARCh samples expressed with increasing induction of the Sec biosynthetic enzymes, using 0/20/80/160 ng/mL aTc respectively. The ion peaks correspond to intact SeARCh (27.9 kDa) and two cleaved sub-sections: amino acids 1-80 (8.9 kDa) and amino acids 81-249 (18.9 kDa). (c) Sampling of parsed spectra resulting from sliding window average applied from 2.5-9.75 min with an averaging window of 0.23 min and single scan offset. Retention times indicate the windows from which shown spectra were generated. (d) Deconvolution of spectra in c. Highlighted regions indicate expected mass of SeARCh1-80 (8878.36 Da), SeARCh81-249 (161S 18965.60; 161U 19027.58), and SeARCh1-249.

Initial mass spectrometry measurements using the purified SeARCh reporter revealed several unusual phenomena which affected the accurate quantification of the relevant protein species. Firstly, consistent with previous reports on the mKate lineage of fluorescent proteins, we observed a spontaneous internal peptide bond cleavage event, evidenced by SDS-PAGE (**Figure S10**)(35). This cleavage yields an N-terminal fragment (residues 1–80) and a C-terminal fragment (residues 81–249), the latter of which contains the UAG codon. Fragment identities were validated with top-down MS2 (**Figure S11, S12**). While we found no evidence that this fragmentation has any impact on fluorescence, it introduced significant complexity to the analysis of the mass spectra, in part due to the highly efficient ionization of the smaller N-terminal fragment. The resulting mixture of Ser- and Sec- containing species, both C-terminal fragments and intact proteins, each with distinct ionization efficiencies, precluded simple quantification. Secondly, our data suggests that the incorporation of Sec may further promote this cleavage event (**Figure 5b**). Specifically, Ser-encoding species appeared to have a lower degree of fragmentation compared to the Sec-containing species. Finally, we observed that as samples approached high Sec:Ser ratios (>80% Sec protein), C-terminal fragments and intact species became increasingly difficult to distinguish and identify (**Figure S8, Figure S13**). This is further compounded by the broad natural isotopic distribution of selenium, and collectively these factors necessitated the development of custom mass spectrometry methods to ensure accurate and reproducible quantification.

Two key steps were taken to address these challenges; liquid-chromatography was used to separate the different species, and peak intensities were summed from smaller windows within the chromatogram rather than averaging across each entire eluting chromatographic peak to preserve masses corresponding to relevant, but low-abundance species (**Figure 5c-d**). In particular, these changes improved the detection of Ser-containing species in samples with high Sec:Ser incorporation ratios, which occur a low abundance but are critical to precise determination of the incorporation fidelity. We further evaluated different data processing approaches to better quantify these rare species and found that an optimized sliding window approach provided the best ability to discriminate between Sec- and Ser-containing forms of the C-terminal fragment as we approached high (∼90%) selenocysteine incorporation fidelity (**Figure S14**). A detailed mass spectrometry workflow is provided in the Methods section.

Leveraging the information gathered using the SeARCh reporter, we sought to define a new minimal Sec biosynthetic pathway and bioprocess conditions that maximize the Sec incorporation efficiency. Using B95.ΔA A*selABCD* cells transformed with our optimal Sec biosynthesis plasmid (**Figure 4d-e, SI Table 1**), shake flask expression with an initial dilution of 1/200 and shorter induction times yielded 5.6 mg/L of purified SeARCh protein with 89-91% Sec incorporation fidelity (**Figure S14c, f**).

## Discussion

The development of successive generations of genetic reporters for selenocysteine has preceded major advances in our understanding of selenocysteine and its biosynthesis and our ability to synthesize new selenoproteins. However, the limitation of these reporters of solely providing information on the magnitude of selenocysteine incorporation makes them ill-suited for solving the current challenge of Ser misincorporation. While analytical techniques such as mass spectrometry can precisely quantify the incorporation ratio of Sec:Ser, these lack the throughput of the molecular methods we might use to tackle this problem. The SeARCh reporter fills this gap and links high throughput, iterative experimentation with *in vivo*, near real-time quantification of the seryl- and selenylation state the tRNA^Sec^ pool. SeARCh leverages the ability of many RFPs to form distinct chromophores with unique spectral properties depending on their protonation state. Here, incorporation of Ser adjacent to the chromophore favours formation of an anionic red chromophore while selenocysteine, which has a far lower pKa, promotes a neutral green chromophore. We show that this two-channel signal is reliably correlated with the Sec:Ser incorporation ratio and is sensitive to subtle changes in the ratio in living cells. Furthermore, SeARCh serves as a reliable proxy for selenoprotein biosynthesis as expressing additional proteins under identical conditions (sfGFP-S205UAG) yields near identical Sec incorporation efficiency, making it a broadly applicable tool for optimizing expression.

By enabling simultaneous monitoring of competing Sec and Ser incorporation using a single protein with an easily measured, quantitative output, the SeARCh reporter can serve as a universal method to standardize the metrology of Sec biosynthesis and incorporation. This addresses a critical challenge in the field where there are often large differences in reported Sec:Ser incorporation ratios between studies which are difficult to rationalize without the ability to make direct comparisons. Using SeARCh, we have identified some of the factors that likely contribute to this variation. Our results clearly indicate that the level of tRNA and target selenoprotein expression have a large influence on the Sec:Ser incorporation ratio. These are both parameters that are expected to vary between studies; expression is highly sensitive to the genetic context of individual experiments, including the promoter or induction strength, plasmid copy number, and codon usage, as well as the inherent variation in expression between different target proteins. By holding expression of the biosynthetic machinery constant and titrating the expression of our SeARCh reporter via an L-arabinose gradient, we were able to change the incorporation ratio by 2.5-fold. This effect was exacerbated with a strong pET expression context where we observed that Sec biosynthesis conditions that previously resulted in mostly green fluorescence, yielded predominately red fluorescence. Collectively, these observations support the hypothesis that charging tRNA^Sec^ with Ser proceeds much faster than the conversion to selenocysteine and that excess Ser-tRNA^Sec^, resulting from either high tRNA expression or rapid turnover of the tRNA^Sec^ pool, will readily become a substrate for translation. Based on this information, both the protein yield and the Sec:Ser incorporation ratio should be taken into account when interpreting past and future results.

Selenoprotein expression is widely assumed to be constrained by Sec biosynthesis; however, our results here paint a more complex picture of the relationship between biosynthesis and the ultimate Sec:Ser ratio measured. We were able to synthesize the SeARCh reporter at yields exceeding 90 mg/L and nearly 80% Sec incorporation fidelity under a variety of conditions, but extensive optimization of the biosynthesis pathway was required to increase the Sec content to 90%. This was primarily achieved through three modifications; increasing the translation rate of SelA, the addition of a new biosynthetic step catalyzed by *Mk*PSTK and shortening the induction period to 6 h. In particular, *Mk*PSTK expression directly correlated with decreased Ser incorporation and makes this hybrid biosynthetic route a compelling alternative to the minimal biosynthetic pathway. The exclusion of Ser is likely a result of the inherent selectivity of the EF-Tu binding pocket. The negatively charged residues Glu216 and Asp217 create an unfavorable environment for negatively charged substrates which for canonical amino acids is offset by stronger interactions with the tRNA(36, 37). Under physiological conditions, the phosphoserine intermediate (SerP-tRNA^SecUXY^) generated by *Mk*PSTK carries a greater negative charge than any canonical amino acid, rendering it a poor substrate for EF-Tu(38). This is supported by the complete absence of any masses associated with SerP incorporation observed by mass spectrometry when *Mk*PSTK was co-expressed with SeARCh. We confirmed that Sep-tRNA^Sec^ was being generated under our expression conditions by showing that SepSecS could complement SelA deletion.

However, these modifications were also associated with a decrease in protein yield. This can be partly explained by a reduction in biomass due to toxicity of highly active Sec biosynthesis but may also be indicative of sequestration of tRNA^Sec^ by SelA and possibly PSTK, which may functionally mirror the effects of reduced tRNA expression. SelA has been previously suspected of impeding selenoprotein production at high expression via tRNA sequestration, and PSTK has been determined to bind both Ser-tRNA^Sec^ and unacylated tRNA^Sec^ with high affinity. The cause of MkPSTK toxicity is currently being investigated but may be due to spurious phosphorylation of host Ser-tRNA^Ser^. Engineered MkPSTK variants which are optimized for interaction with bacterial tRNA^Sec^ may resolve both of these problems(39).

One of the most significant gaps in our understanding of selenocysteine remains the initial steps of Se assimilation, including the identity of the native selenium donor for SelD which may be the interface between inorganic selenium metabolism and Sec biosynthesis. Using SeARCh, we show that the thioredoxin system is not strictly required for Sec biosynthesis nor activity of *E. coli* FDH_H_ which contradicts prior reports. The reliance on reporters that are dependent on other cellular processes may conflate disruptions in reporter activity with disruptions in Sec biosynthesis. Indeed, FDH_H_ requires both iron-sulfur cluster assembly and synthesis of a molybdopterin cofactor, either of which may be susceptible to defects in redox metabolism in different strain backgrounds or thioredoxin specifically(40). This makes the enzyme poorly suited as a reporter for reductive assimilation of inorganic selenium which is intertwined with sulfur and redox metabolism. Individual disruption of several genes relating to these processes did not identify any new candidates upstream of the terminal biosynthetic machinery that affected Sec biosynthesis, possibly due to significant redundancy. Higher-order genetic knockouts may be needed to more thoroughly investigate these steps, but this is complicated by the fact that these processes are essential for cell viability. Disruption of either the thioredoxin or glutathione systems in this work had no observable effect, but disruption of both results in significant growth defects which would be difficult to separate from specific impacts on Sec biosynthesis. We anticipate that the two clearly distinct emission maxima or SeARCh (>100 nm apart) will enable high-throughput functional genomics approaches to more broadly identify new genes which influence Sec biosynthesis.

Mass spectrometry is the definitive standard for reporting Sec:Ser incorporation ratios in recombinant selenoproteins. However, there are several confounding factors that can complicate mass spectrometric analysis. Selenium itself has an unusual isotope distribution with six naturally occurring stable isotopes (^74^Se, 0.89%; ^76^Se, 9.37%; ^77^Se, 7.63%; ^78^Se, 23.77%; ^80^Se, 49.61%; ^82^Se, 8.73%), several of which are highly abundant(41, 42). This isotope distribution in combination with the heterogenous expression of both Sec- and Ser-containing protein species can yield overlapping ion peaks that require careful analysis to parse the identity and quantity of each species. In the case of the SeARCh reporter, this is further complicated by the unique backbone cleavage event which itself appears to be differentially affected by the incorporation of Sec or Ser. These issues can obscure rarer species that may be difficult to resolve from the background or may be erroneously eliminated during data processing. This is particularly a problem as we seek to close the gap towards selenoproteins with near quantitative Sec incorporation and the competing Ser species become increasingly rare. To ensure the most accurate accounting of all protein species relevant for quantification of Sec incorporation we compared multiple data processing techniques and optimizations thereof. An optimized sliding window spectral averaging method provided the most complete account of all expected species, including the rare Ser species lost as a result of other spectral averaging methods, including parsed windows or a broad single average. A detailed report on these steps is provided in the methods to facilitate further characterization of the SeARCh reporter and accommodate its unusual peptide backbone cleavage event.

Here we describe the SeARCh reporter which provides unique spectral outputs corresponding to Sec or Ser and demonstrate its broad utility for measuring and optimizing Sec biosynthesis. SeARCh offer considerable advantages over other genetic reporters which are incapable of providing information on Ser misincorporation and may be influenced by other changes in cellular metabolism. Strategies to improve the reporter by increasing the brightness of the green chromophore and reducing spontaneous backbone cleavage are being investigated. Informed by SeARCh, we have developed several improvements to the minimal Sec biosynthesis pathway and identified several phenomena which likely contribute to variation in the Sec incorporation fidelity reported in the literature. While there are still significant knowledge gaps around Sec biosynthesis, we anticipate that these new tools will enable more precise and reproducible engineering efforts in the future.

## Methods

### Molecular Cloning

Plasmids used in this work were constructed either using Gibson Assembly or a Golden Gate based hierarchical modular cloning system. Gibson assemblies were incubated at 50 °C for at least 90 minutes. Golden Gate cloning was performed by combining 25 ng of each plasmid containing a modular DNA fragment and subjected to 30 cycles of digestion and ligation (37 °C for 1 minute followed by 16 °C for 2 minutes). All assemblies were transformed into either *E. coli* DH10B or DB3.1 using standard methods.

### Fluorescence Assays

All SeARCh fluorescence assays were performed in *E. coli* strain B95ΔA Δ*selABCD* Δ*fabR* in which the suf operon had been repaired. The assays were performed using the following media compositions: M9 + yeast extract (2.5 g.L^-1^) + 0.5% glycerol, M9 + yeast extract (0.5 g.L^-1^) + 0.5% glycerol, Terrific Broth (TB) + 0.5% glycerol, or ½ Terrific Broth + 0.5% glycerol. Transformants were selected and cultured in 1 mL volumes in a 96-well grow block with 900 rpm agitation on a 1.5 mm orbit. For assays containing a pET vector, the cells were grown in the presence of 2% glucose. Following incubation overnight at 37 °C, cells were diluted 1:33 into fresh media with Na_2_SeO_3_ and treated as follows. For assays where a component or components of the biosynthetic pathway were induced, the cells were incubated for 1 hour after dilution before aTc (final concentration of 160 ng/mL unless otherwise specified) was added. After an additional hour of incubation, either L-arabinose (final concentration of 0.2% unless otherwise specified) or IPTG was added. The cultures were incubated for either 4 or 16 hours at 37 °C before the cells were harvested by centrifuged at 3500 x g at 4 °C for 10 minutes. Supernatant was decanted and the cells were resuspended in 1 mL of PBS before being transferred to a 96-well assay plate. Each well was measured for absorbance at 600 nm (OD_600_), green fluorescence (445 nm *ex*, 525 nm *em*), and red fluorescence (585 nm *ex*, 635 nm *em*).

### Flask Protein Expression

Cells were treated similarly as described above with the following modifications. Overnight cultures were inoculated in test tubes and were diluted 1:200 into 200 mL of fresh media in a 500 mL baffled flask. The cultures were incubated at 37 °C with shaking until an OD_600_ of 0.2-0.3 was reached before aTc was added at a final concentration of 160 ng/mL (unless otherwise specified). After an additional hour IPTG was added to a final concentration of 50 μM (unless otherwise specified) and cultured for 6 or 16 hours. A 1 mL aliquot was centrifuged to perform fluorescence measurements as described above.

### Protein Purification

Cells expressing SeARCh or sfGFP using a pET vector system with an added 6xHis tag with a thrombin site on the N-terminus were harvested by centrifugation and resuspended in IMAC Resuspension Buffer (50 mM NaH_2_PO_4_, 100 mM NaCl, 1 mM EDTA, pH 8.0) and lysozyme was added at a concentration of 3 mg.mL^-1^. The cells were incubated at 37 °C for 30 minutes with shaking and then lysed by sonication. The lysate was clarified by centrifugation and the EDTA quenched by addition of MgCl_2_ at a final concentration of 10 mM. The supernatant was purified using Ni-NTA resin and eluted with IMAC elution buffer (50 mM NaH_2_PO_4_, 300 mM NaCl, 300 mM imidazole, pH 8.0). Proteins were dialyzed into a sodium phosphate buffer (20 mM NaH_2_PO_4,_ 50 mM NaCl, and pH 7.4) for storage and further characterization.

### CRISPR-Associated Transposase (CAST) Gene Disruptions

Electrocompetent B95.ΔA.Δ*selABCD* cells were electroporated with a plasmid encoding the CAST machinery and a crRNA guide targeting a site within 50 base pairs upstream of the gene of interest. Following recovery in SOC medium for 1 hour at 37 °C, cells were plated on selective agar and incubated overnight at 30 °C. Resulting colonies were screened for targeted disruption via colony PCR using primers flanking the disruption site. For functional characterization, these engineered strains were subsequently transformed with the SeARCh reporter and Sec biosynthetic machinery according to the protocols described above.

### Mass Spectrometry of paired SeARCh and sfGFP samples

GFP variants were reduced with 5 mM dithiothreitol for 30 minutes at 55°C and alkylated with 15 mM iodoacetamide for 30 minutes at room temperature in the dark to cleave cysteine – selenocysteine bonds. Both GFP and SeARCh variants were diluted to 10 μM and desalted using a P6 Micro Bio-Spin columns into 50:50 methanol:water with 0.1% formic acid for mass spectrometry analysis. Protein solutions were analyzed on a Thermo Orbitrap Eclipse mass spectrometer (Thermo Fisher, San Jose, CA). Proteins solutions were introduced with nanoelectrospray ionization using a Au/Pd coated borosilicate static tip pulled in-house (OD 1.2 mm, ID 0.69 mm) and a 1.4 kV applied voltage. Spectra were collected at 240 K resolving power using 200 averages, with full profile mode, and deconvoluted with Xtract in Freestyle with a S/N of 10. Semi-quantitative analysis was done using the peak heights of the two variants based on the deconvoluted MS1 spectra in Xtract. For the SeARCh protein, the peaks for the intact protein and the C-terminal fragment containing amino acids 81-249 were added together for quantitative analysis.

### Liquid Chromatography Mass Spectrometry Analysis of Intact SeARCh

Intact mass analysis experiments were performed on a Thermo Scientific Vanquish Neo chromatography system coupled to a Thermo Scientific Orbitrap Ascend equipped with a nanoflex NG source. Proteins were subjected to acetone precipitation then resuspended in water with 0.1% FA to a final concentration of 200 ng/uL and injected onto a Waters NanoEase m/z BEH C4 column (300 A pore size, 1.7 um particle size, 150um I.D. x 100mm). Proteins were eluted at a constant flow rate of 2 uL/min over a multi-segment gradient: 5-27% ACN in 2 min; 27-33% ACN in 10 min; 33-97% ACN in 1 min. MS1 spectra were acquired from *m/z* 500-2000 at a resolving power of 120000 with 1 microscan.

For tandem mass spectrometry experiments, MS1 scans were performed identically as for intact mass analysis (**Figure S11**). Precursor ions were automatically selected for fragmentation with an isolation width of 1.6 *m/z*, a charge range filter of +18 to +30 was applied for selection. Proteins were activated using EThcD (reaction time 5 ms; ETD reagent target 7.0e5; maximum reagent injection time 200ms; HCD collision energy 15%) or UVPD (activation time 50 ms). MS2 scans were acquired in the Orbitrap analyzer with 25 microscans at a resolving power of 120000. For experiments utilizing a single fragmentation method, cycle time was 15 s; for experiments using both UVPD and EThcD fragmentation methods, cycle time was 30s.

### Spectral Deconvolution

MS1 data was deconvoluted using Thermo Scientific Biopharma Finder 5.1 in Intact Mass Analysis mode or UniChrom within UniDec v 8.0.3(43). For analysis using Biopharma Finder, spectra were averaged using the sliding window method (target average spectrum width: 1.12 min; scan offset: 1) and deconvoluted using the Xtract deconvolution algorithm with the following parameters: Output mass: 1000-100000 Da; S/N threshold: 3; Relative abundance threshold: 0%; Charge range: 5-60; Isotope table: protein. Sum intensity of identified species in the deconvoluted spectra was used for relative quantification of selenocysteine incorporation efficiency. For UniChrom analyses, spectra were partitioned into windows ranging from 0.23-2.5 min wide. Uni Dec deconvolution was performed with a charge state range of 5-50, peak FWHM equal to 1, gaussian peak shape, with 50 maximum iterations. Peaks were picked and extracted with a peak picking range of 15 Da and a threshold of 0.1.

### Tandem Mass Spectrometry Data Analysis

MS2 data was processed with Prosight PD 4.3 (Proteinaceous) against a custom database containing all expected proteoforms of the recombinant protein sequence in an error-tolerant fashion to account for expected error in the deconvoluted mass of selenocysteine containing proteins (**Figure S12**). Mass tolerance for fragment matching was set to 20 ppm. Proteoform spectrum matches (PrSMs) and proteoforms were validated with an FDR cutoff of 1%.

### Replicates and Statistics

All experiments were performed in biological and technical triplicate unless specifically indicated. Fluorescence data is plotted as the mean ± std, unless specifically stated otherwise.

## Data Availability

The genetic materials and data that support the findings of this study are available from the corresponding author upon reasonable request.

## Acknowledgements

The authors thank the Welch Foundation for their support (C-2167-20230405 to Ross Thyer and F-1155 to Jennifer S. Brodbelt). Jennifer S. Brodbelt acknowledges support from National Institutes of Health (R35GM139658). Devon L. Kulhanek is grateful for the support of the Graduate Research Fellowship Program (National Science Foundation). This work was completed in part using instrumentation of the Shared Equipment Authority at Rice University.

## Author information

Authors and Affiliations

Systems, Synthetic, and Physical Biology, Rice University, Houston, USA

Andrew Gilmour and Zach Jansen

Department of Bioengineering, Rice University, Houston, USA

Qiyao Wei

Department of Chemistry, The University of Texas at Austin, Austin, USA

Jessica Hellinger and Jennifer S. Brodbelt

Department of Chemical and Biomolecular Engineering, Rice University, Houston, USA

Devon L. Kulhanek and Ross Thyer

Shared Equipment Authority, Rice University, Houston, USA

Katelyn M. Baumer

## Author Contributions

**Andrew Gilmour:** Conceptualization; Methodology; Data curation; Investigation; Formal analysis; Writing – original draft, review and editing. **Qiyao Wei:** Conceptualization; Data curation; Formal analysis; Writing – review and editing. **Jessica Hellinger:** Investigation; Formal analysis; Writing – review and editing. **Devon Kulhanek:** Methodology; Writing – review and editing. **Zachary Jansen:** Methodology; Writing – review and editing. **Katelyn M. Baumer:** Investigation; Formal analysis; Writing – review and editing. **Jennifer S. Brodbelt:** Supervision; Writing – review and editing. **Ross Thyer:** Conceptualization; Supervision; Methodology; Writing – review and editing.

## Ethics declaration

### Competing interests

The authors declare no competing interests.

## Supplementary information

Supplementary Figs. 1–14, and Table 1.

## Notes

### Competing Interest Statement

The authors have declared no competing interest.

