## Supplementary Information for "A Quantitative Two-Channel Genetic Reporter for Selenocysteine Biosynthesis and Incorporation"

This file contains:

Supplementary Figures S1-S14

Supplementary Table 1

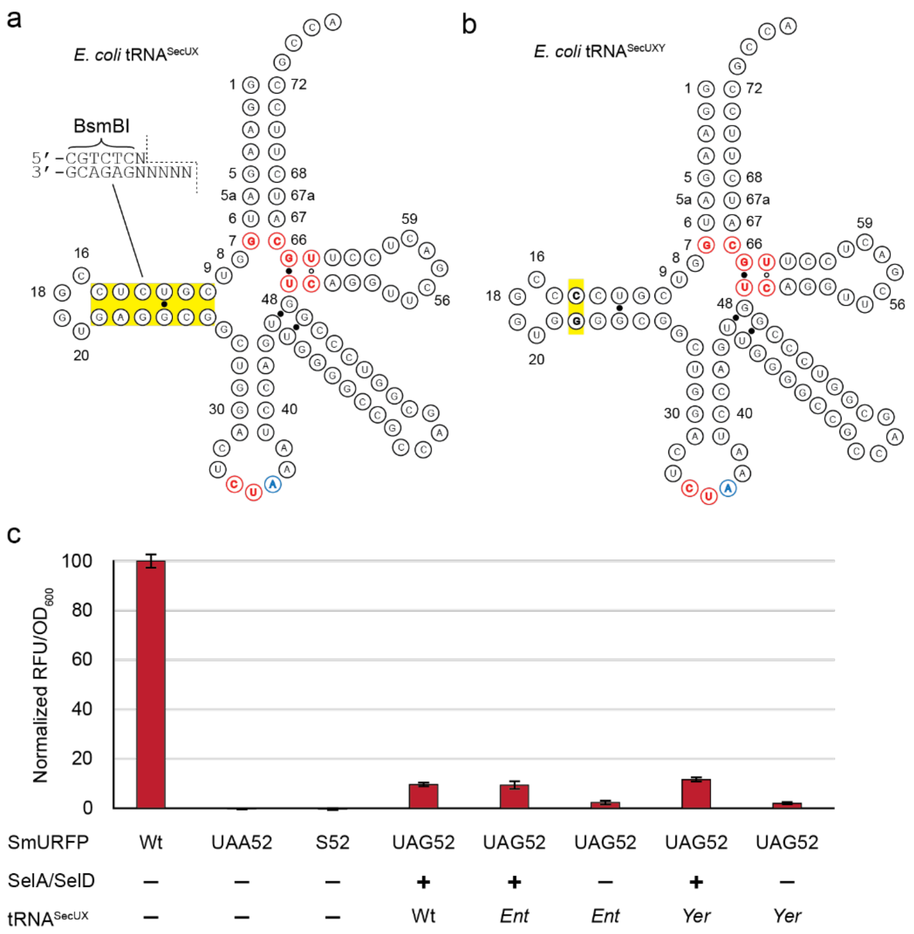


**Figure S1. Domestication of tRNA$^{SecUX}$ for Esp3I (BsmBI) compatibility.** (a) Structure of tRNA$^{SecUX}$ with D-arm sequence containing Esp3I cut-site highlighted in yellow. Red bases at base of acceptor stem and T-arm represent mutation introduced previously to allow for interaction with *E. coli* EF-Tu. Colored bases at the bottom of the structure represent the anti-codon for UAG. (b) Structure of tRNA$^{SecUXY}$ with domestication mutations ($C_{14}$-$G_{22}$) highlighted in yellow (c) Seleno-SmURFP fluorescence assay validation of tRNA$^{SecUX}$ mutations. Incorporation of either a cysteine (wt) or selenocysteine (UAG52) is required for ligation of the biliverdin chromophore and fluorescence ($\lambda_{ex}$: 635 nm, $\lambda_{em}$: 685 nm) while termination (UAA52) or serine (S52) renders the protein non-fluorescent. Both versions of tRNA$^{SecUX}$ that either harbored mutations from *Yersinia enterolitica* or *Enterobacter sp.638* showed identical activity to wild-type in the presence of SelA and SelD. (n=3, ±std).

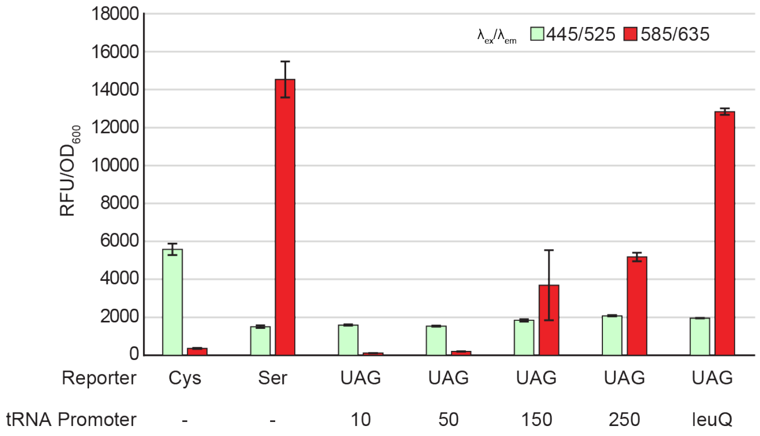


**Figure S2. SeARCh pET expression assay.** Fluorescence assay of SeARCh reporter expressed with a strong T7 promoter on a pET vector using a subset of $tRNA^{SecUXY}$ constitutive promoter strengths. A systematic increase in red signal ($\lambda_{ex}$: 585 nm, $\lambda_{em}$: 635 nm) is observed as tRNA promoter strength increases, indicating a correlation between $tRNA^{SecUXY}$ abundance and serine incorporation. (n=3, ±std).

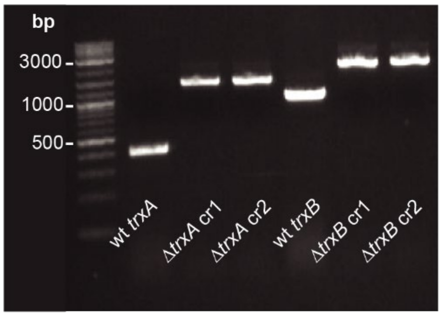


**Figure S3. Validation of CAST-mediated host gene disruptions.** PCR analysis of *trxA* and *trxB* loci confirms successful integration of the CAST transposon. Disruption yields an expected insertion of ~1000bp. Disruptions were generated using two independent crRNA guides (cr1 and cr2) for each gene.

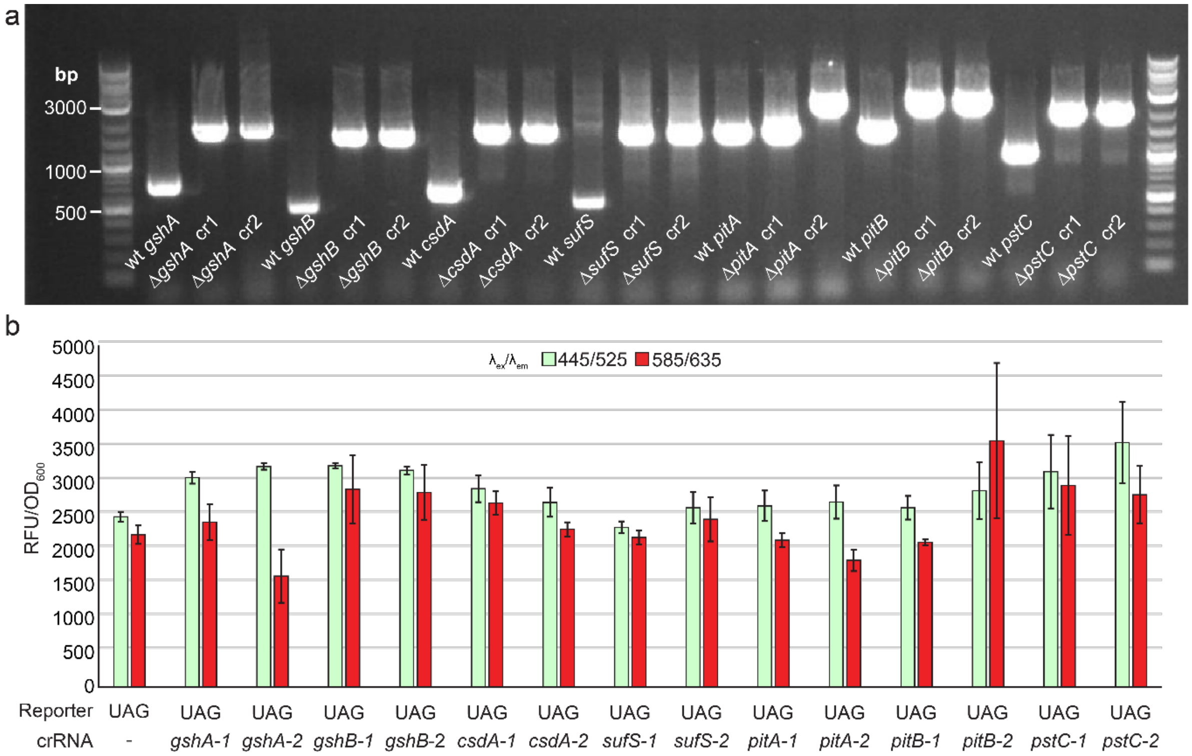


**Figure S4. Gene disruptions do not impact Sec incorporation.** (a) PCR analysis confirming transposon integration. Disruption yields an expected insertion of ~1000bp. Two independent crRNA guides (cr1 and cr2) were used for each gene. Note: *pitA* cr1 did not yield successful integration and is represented by a wild-type amplicon. **(b)** Fluorescence assay using the pET-SeARCh reporter and Sec biosynthesis machinery to evaluate if gene disruptions modulated Sec incorporation. Note: *pitA-1* represents wild-type. (n=3, ±std).

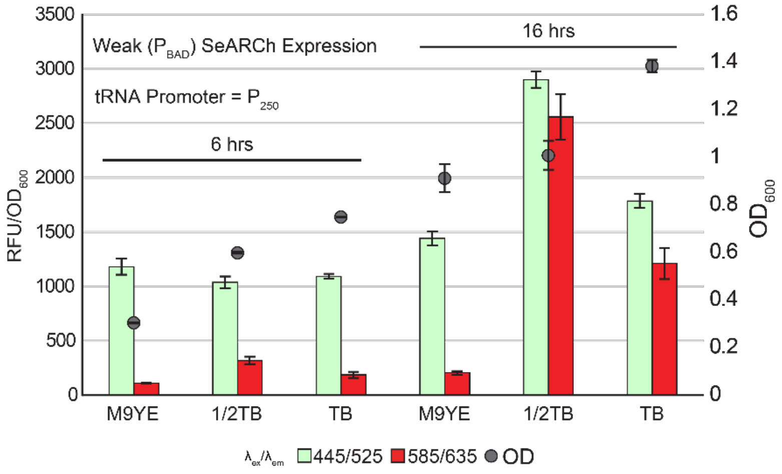


**Figure S5. Optimization of media composition and induction timing.** SeARCh reporter fluorescence and cell growth (grey circles, $OD_{600}$) were compared between three different media (M9YE, 1/2 TB, and TB) and two induction durations (6 and 16 hours). Longer induction durations are accompanied by an increase in both $OD_{600}$ and overall fluorescence. Richer media compositions correlate with higher $OD_{600}$ across both induction durations display increased red fluorescence at longer durations. (n=3, ±std).

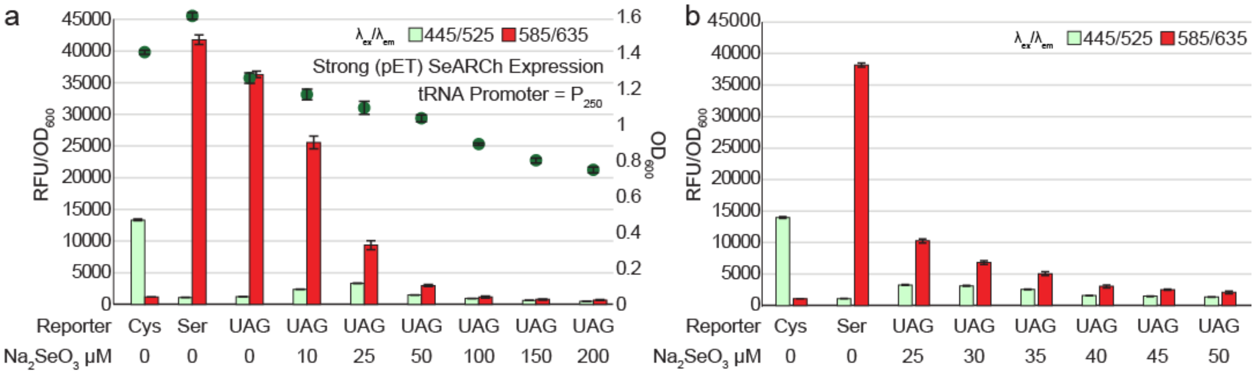


**Figure S6. Optimization of exogenous selenium supplementation.** (a) SeARCh reporter fluorescence and cell growth (grey circles, $OD_{600}$) were compared across a gradient of 0-200 μM $Na_2SeO_3$. A strong correlation between increasing $Na_2SeO_3$ and decreasing red signal ($\lambda_{ex}$: 585 nm, $\lambda_{em}$: 635 nm) and $OD_{600}$ is observed. (b) High resolution titration of exogenous [Se] over a gradient of 25-50 μM $Na_2SeO_3$. 30 μM $Na_2SeO_3$ was determined to be the threshold before protein expression and $OD_{600}$ decrease. (n=3, ±std).

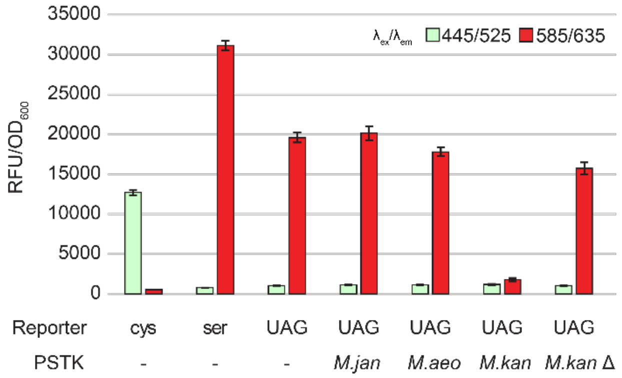


**Figure S7. Comparative analysis of Archaeal PSTK activity toward tRNA$^{SecUXY}$.** Three distinct archaeal PSTK homologs were independently transformed into B95.ΔA Δ*selABCD* harboring the pET-SeARCh reporter and a constitutively expressed Sec biosynthetic pathway. A truncated variant (Δ*Mkan*) consisting of the N-terminal 89-residues, was included as a negative control. The reduction in red signal ($\lambda_{ex}$: 585 nm, $\lambda_{em}$: 635 nm) suggests *Mk*PSTK successfully phosphorylates the tRNA$^{SecUXY}$, precluding it from entering translation. (n=3, ±std).

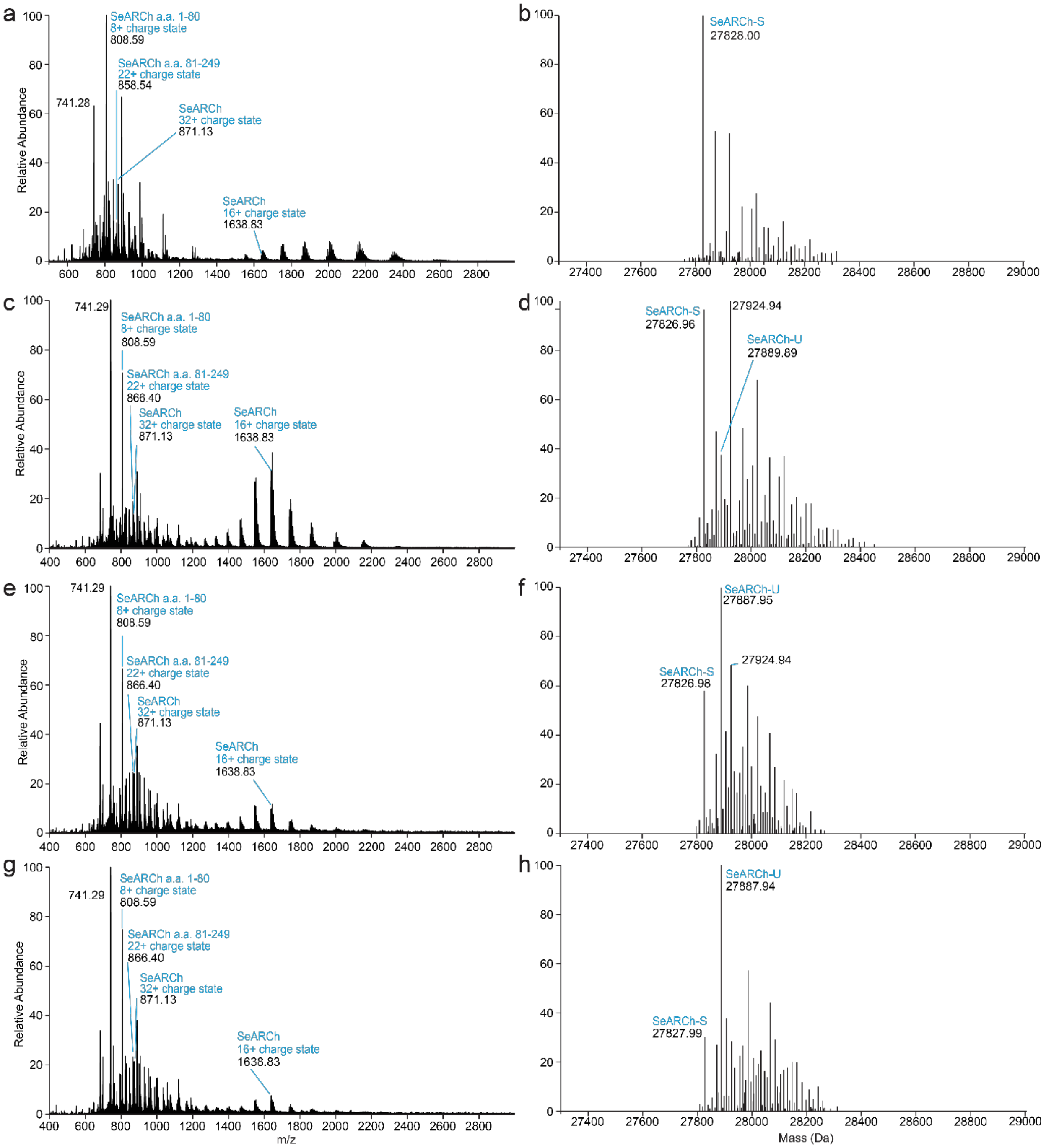


**Figure S8. Matched MS1 (left side) and deconvoluted (right side) spectra of SeARCh.** Samples represent SeARCh protein expressed in conjunction with increasing aTc concentrations to induce higher expression of the Sec biosynthetic enzymes. Sec biosynthesis induction: 0 ng/mL aTc (a-b), 20 ng/mL aTc (c-d), 80 ng/mL aTc (e-f), and 160 ng/mL aTc (g-h). The deconvoluted spectra on the right only show the mass region corresponding to the intact proteins. The additional unlabeled peaks correspond to adducts.

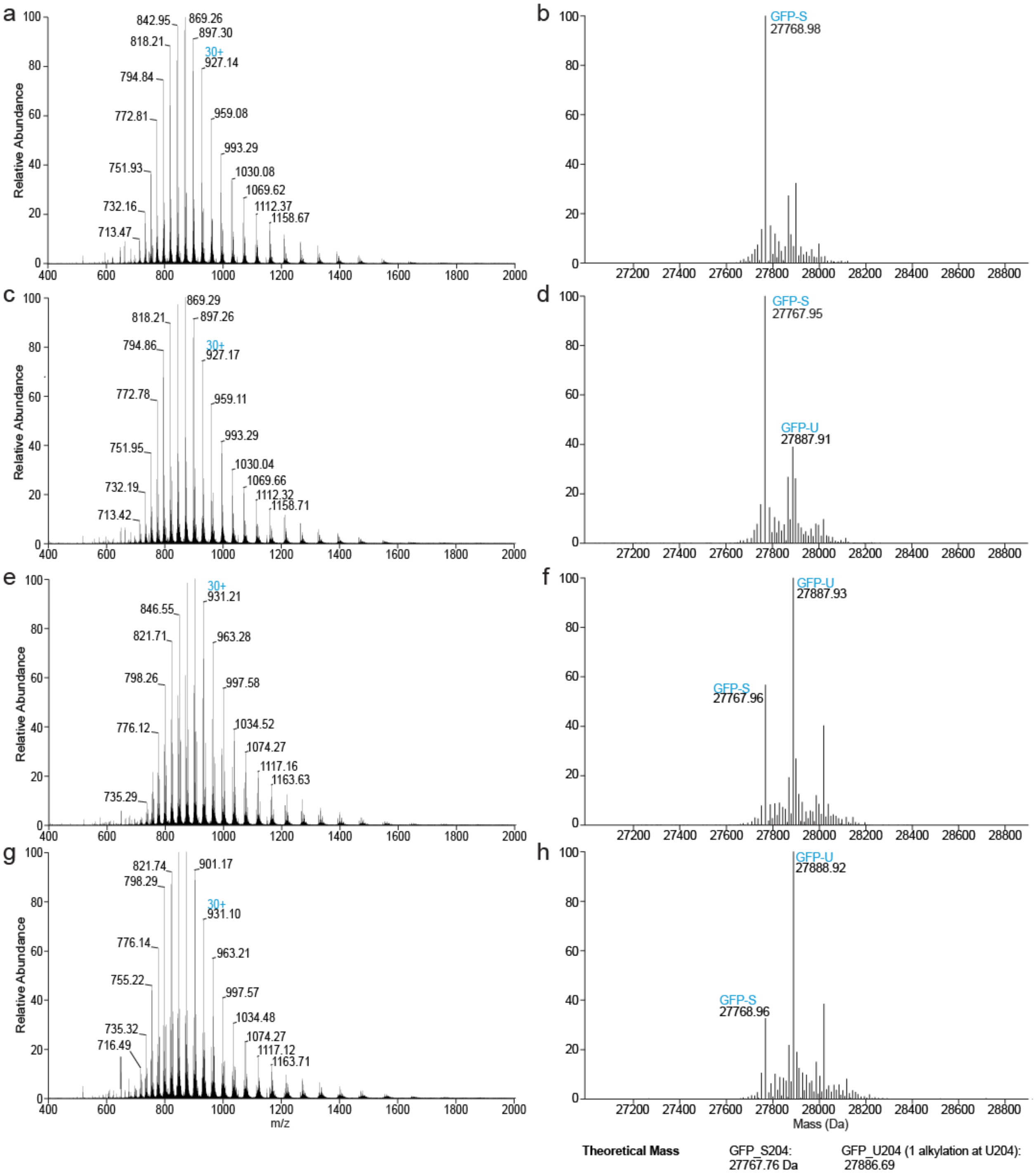


**Figure S9. Matched MS1 (left side) and deconvoluted (right side) spectra of sfGFP.** Samples represent sfGFP protein expressed in conjunction with increasing aTc concentrations to induce higher expression of the Sec biosynthetic enzymes. Sec biosynthesis induction: 0 ng/mL aTc (a-b), 20 ng/mL aTc (c-d), 80 ng/mL aTc (e-f), and 160 ng/mL aTc (g-h). The deconvoluted spectra on the right only show the mass region corresponding to the intact proteins.

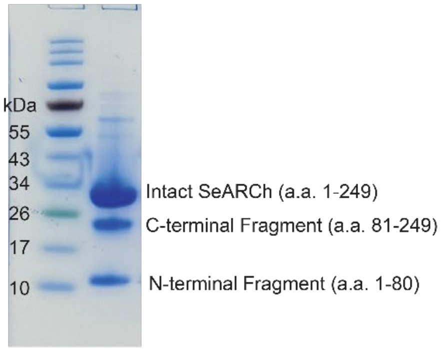


**Figure S10. SDS-PAGE of purified SeARCh displaying fragmentation.** SeARCh encoding an N-terminal 6xHis tag was purified and separated on a 4-12% SDS-PAGE gel. Three bands corresponding to the intact protein (residues 1-249), the C-terminal fragment (residues 81-249), and the N-terminal fragment (residues 1-80) are the predominant species.

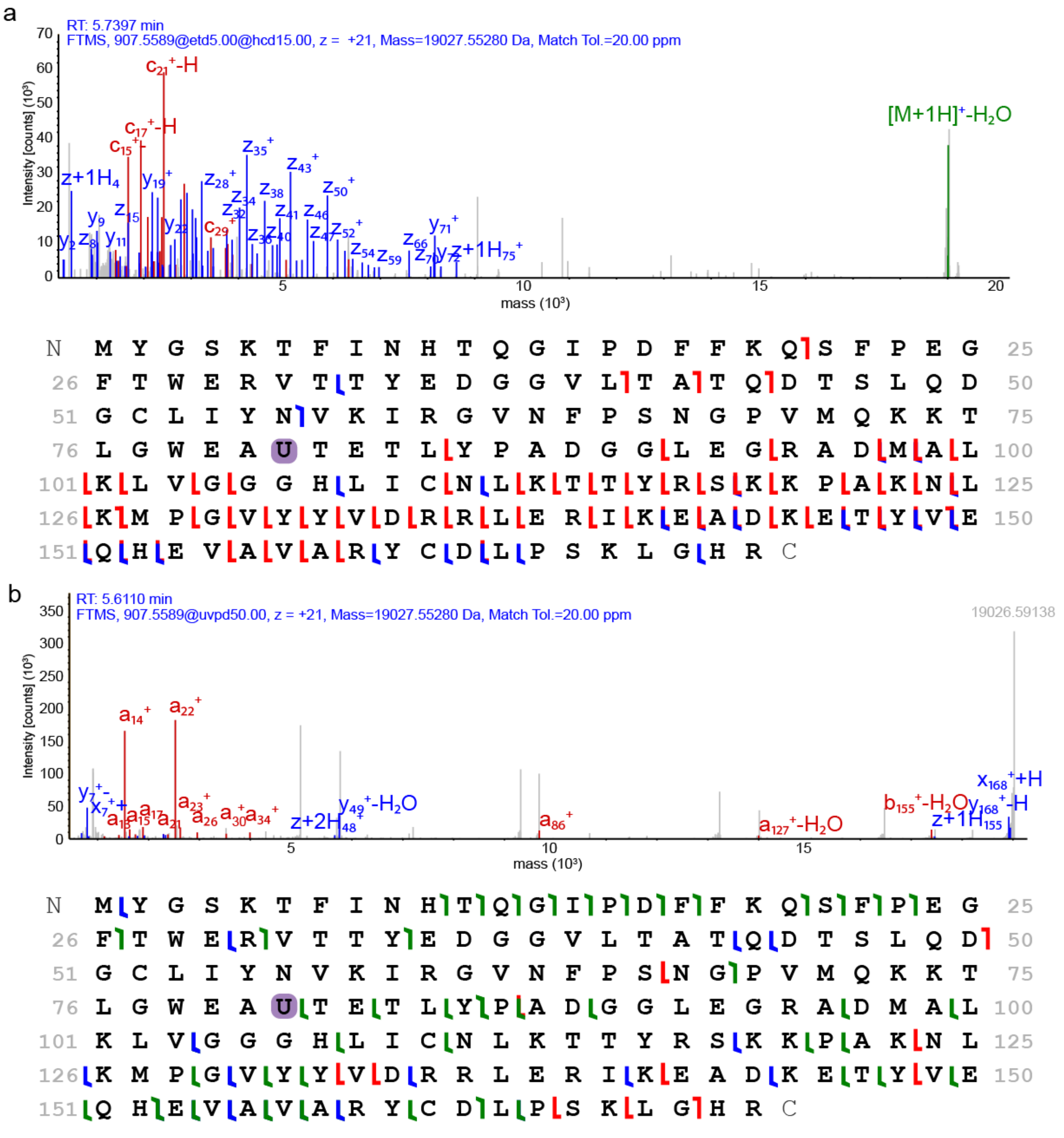


**Figure S11. Top-down fragmentation confirms identity of SeARCh$_{81\text{-}249}$.** Tandem mass spectrometry was used to confirm identity of SeARCh cleavage products. Proteins were fragmented using (A) EThcD or (B) UVPD. Fragment matches were assigned with a mass tolerance of 20 ppm. All PrSMs and proteoforms were assigned in ProSight PD 4.3 with an FDR cutoff of 1%.

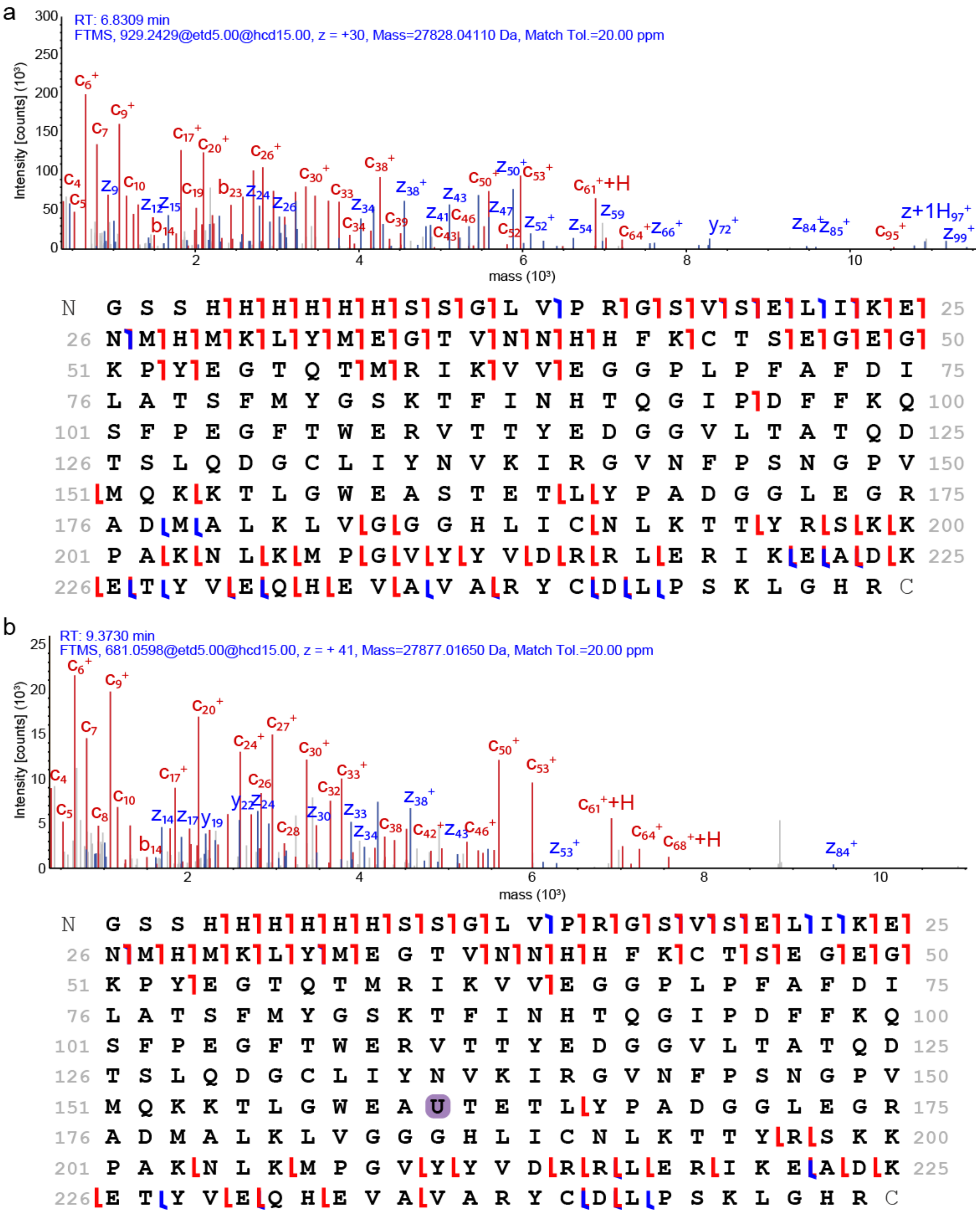


**Figure S12. Top-down fragmentation confirms selenocysteine incorporation in full-length SeARCh.** MS2 spectra and corresponding fragment maps for (a) serine containing, full length SeARCh or (b) selenocysteine containing, full length SeARCh. Fragment matches were assigned with a mass tolerance of 20 ppm. All PrSMs and proteoforms were assigned in ProSight PD 4.3 with an FDR cutoff of 1%.

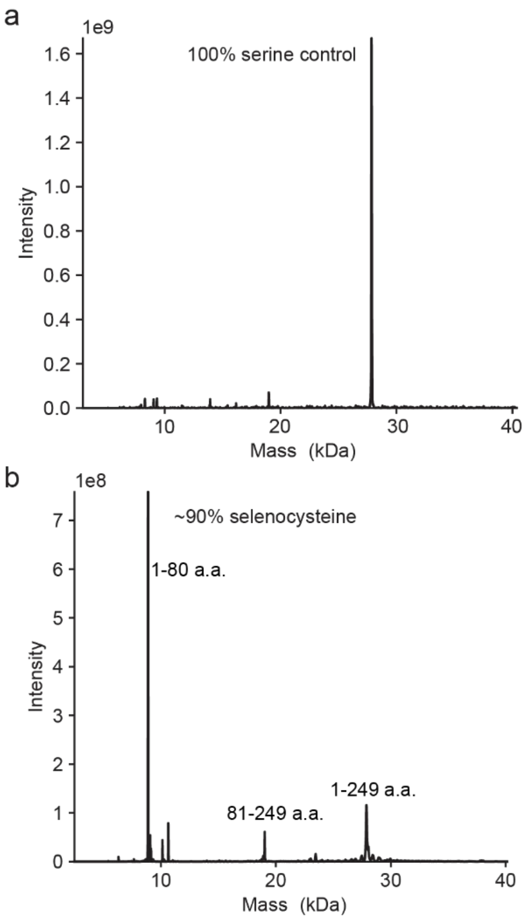


**Figure S13. Comparison of fragmentation of serine and selenocysteine forms of the SeARCh reporter in the deconvoluted mass spectra based on MS1 analysis of the protein samples.** (a) Deconvoluted mass spectrum of a serine-only control of the SeARCh reporter showing predominantly one peak consistent with the expected mass of the full-length protein. (b) Deconvoluted mass spectrum of a sample with ~90% selenocysteine showing the presence of the two fragments corresponding to the first 1-80 and last 81-249 amino acids as well as a full-length intact protein (1-249 a.a.).

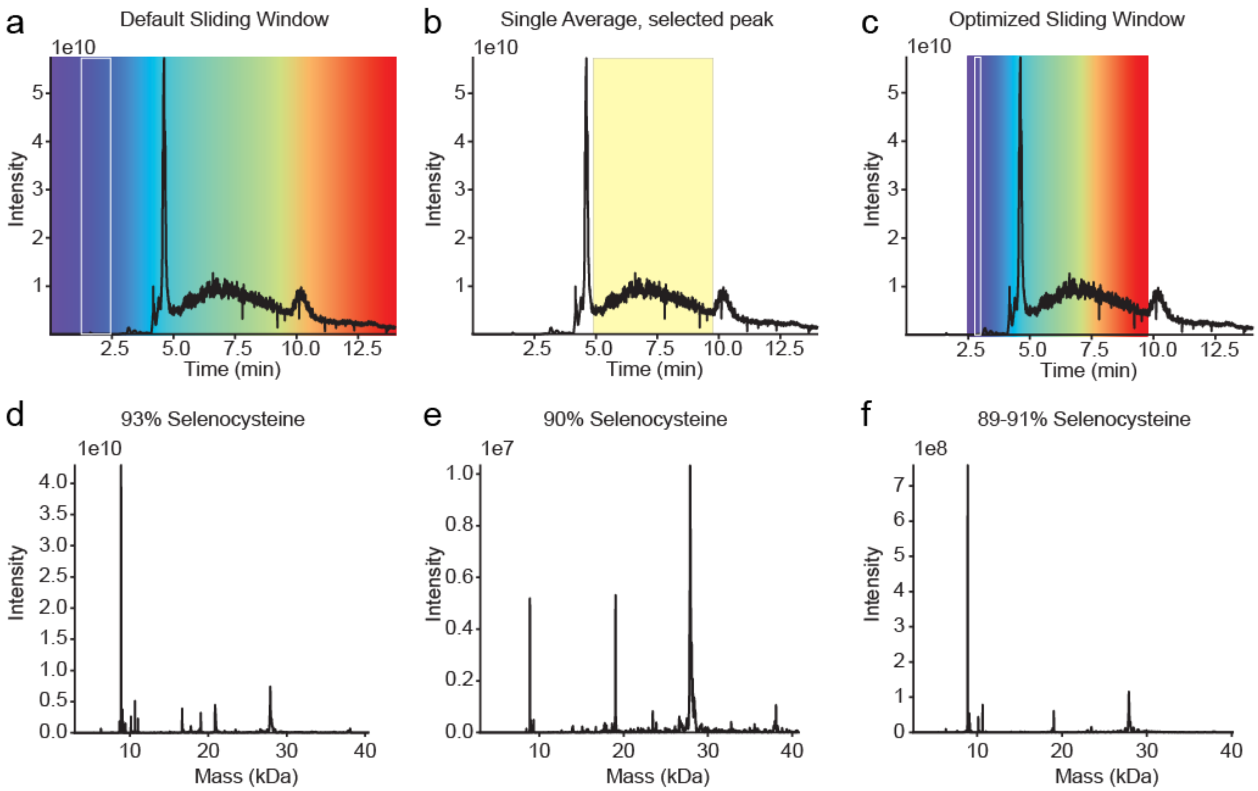


**Figure S14. SeARCh reporter mass spectrometry processing comparison.** Total ion current chromatograms (a-c) and paired composite deconvolution after spectral averaging (d-f) which results in varying ability to capture rare Ser-containing species. Spectral averaging was performed as follows: (a,d) sliding window width = 1.12 min, scan offset = 1, retention time range 0-15 min; (b,e) a single spectral average ranging from 5-9.75 min; or (c,f) sliding window width = 0.23 min, offset = 1 scan, retention time range 2.5-9.75 min. The width of windows in (a) and (c) are further denoted by the white outline of a single window. Spectra in (d) and (f) represent the summed composite of all sliding windows.

**Table S1:** DNA sequences of SeARCh, tRNA$^{SexUXY}$, *Mk*PSTK, and our final selenocysteine biosynthetic expression vector that contained SelA/D/tRNA$^{SecUXY}$ and *Mk*PSTK. Start codons are underlined and SelA (yellow), SelD (red), tRNA$^{SecUXY}$ (magenta), and *Mk*PSTK (teal) sequences are highlighted in color.

| Coding Sequences |
|---|
| SeARCh (mKate2 A45V S143U) |
| atggtgtccgaattgatcaaggagaacatgcacatgaagctttatatggagggtacagttaacaaccac<br>cactttaaatgtacctcagaaggcgaaggtaaaccttacgaaggtacgcagaccatgcgtattaaggtg<br>gtcgaaggcggcccgttaccattcgcctttgatattctggcgactagttttatgtacggttccaaaact<br>ttcatcaaccatacacagggaatcccggatttcttcaagcagtcgtttcccgaggggtttacgtgggag<br>cgcgtgacgacgtatgaggacggcggggtgttaactgcgacacaggacacgagtctgcaggacggttgt<br>ttgatctacaatgtcaagatccgcggtgttaatttcccctcaaatggacctgttatgcaaaagaaaact<br>ctgggttgggaggcatgtaccgaaacgctttacccagctgatggggcctggagggccgcgcggatatg<br>gcgttaaaactggtaggcggtggacacctgatttgtaaccttaagacgacttatcgcagtaagaaaccc<br>gcgaaaaacctgaaaatgcctggtgtttactatgttgaccgccgtttagagcgcattaaagaagctgat<br>aaggaaacctacgtcgaacaacacgaggtggccgttgctcgctactgtgacttacccagtaaactggga<br>catcgctaa |
| tRNA$^{SecUXY}$ |
| ggaagatggtcgtccccggtggggcggctggactctaaatccagttggggccgccagcggtcccggtca<br>ggttcgactccttgcatcttccgcca |
| *Mk*PSTK |
| atgcgactgctgatactcaccggcccgcccgggtcaggtaaaacgtgtttcgcgcgggagctggctcgg<br>gaactccggcaggaaggctggcgagtggcccatgtggaggcggacgctctccgcggattcctatgggat<br>gagttcgacccgaaactcgagcaagtggcccgggaactcttttttgaagtccgttgaaacgtgtctggac<br>gccgagctagacctggtgatagcggacgacactaattactattccagcatgcggcgtgagttggctctc<br>ctggccttggagaggaaagttccgtgggggatagtgtacctccgtaccggtttagatacgtgcctacgt<br>aggaaccgcgagcgaggtgaacctatcccggaggaggtcgttcgtagaatttatgataggtttgaacca<br>cccgagccggatcggtggtgggagcgtgcgaccttggttctagacgattctcgggtttccgaggaagtt<br>ctcgagttcgtcgagtcgggtctacgcgtcgagaagcccaaaaagcgacgtcggcgcactgacccctcc<br>tccgtgaatgaagtcgacgtacgcacccgtcaggttatgggggagctgatgcgtcgactctccgagaca<br>ggcgcggctacccaagagctagggcgcaagctcagtgaactgcgccgtgagatagtgtcctccgtggag<br>gatccggaaaaggccgtccgggagttccggcgtcggccgaggaggtgatccgggaatgcctgcatgga<br>gatgggtga |
| Sec biosynthetic pathway plasmid – Figure 4e |
| tccctatcagtgatagagattgacatccctatcagtgatagagatactgagcacatcagcaggacgcac<br>tgaccggagagctgtcaccggatgtgctttccggtctgatgagtccgtgaggacgaaacagcctctaca<br>aataattttgtttaagggcccaagttcacttaaaaaggagatcaacaatgaaagcaattttcgtactga<br>aacatcttaatcatgcgccggtgttttaaatatgactactgaaacgcgtttcctgtacagccagctgcc<br>ggctattgaccgcctgctgcgcgactcctcttttctgagcctgcgtgacacctacggtcacacccgcgt<br>ggtagagctgctgcgtcagatgctggatgaagcgcgtgaggtaatccgtggcagccaaccctgccggc<br>ttggtgcgaaaactgggcgcaggaagtagatgcgcgcctgactaaagaagctcagtctgccctgcgtcc<br>ggtaatcaacctgaccggcactgttctgcacaccaacctgggccgtgctctgcaggcagaagccgctgt<br>agaggcggtagctcaggctatgcgctctccggttaccctggaatacgacctggatgacgcaggtcgtgg<br>tcaccgcgatcgtgcgctggcgcaactgctgtgtcgtattactggcgcggaagatgcgtgcatcgttaa<br>taataacgcggcggcagtcctgctgatgctggcagcgaccgcgtccggcaaggaagtggtagtatctcg<br>cggcgaactggttgagattggcggcgcgttccgcatcccggatgttatgcgccaggccggttgcaccct<br>gcatgaagtaggcactaccaaccgtacccacgctaacgattatcgtcaggcagttaacgaaaacaccgc<br>cctgctgatgaaagttcacacgagcaactactccatccagggcttcaccaaagcaatcgacgaagccga<br>actggttgctctgggcaaagaactggacgttccggttgtaaccgatctgggttctggcagcctggtgga<br>cctgtctcagtacggcctgccgaaagaaccgatgccacaggaactgattgcagcgggtgtttctctggt |

gagcttctctggcgacaaactgctgggtggtccgcaggctggtatcattgtaggtaaaaagaaatgat
cgcgcgtctgcagtctcaccctctgaagcgcgctctgcgtgcagacaaaatgaccctggctgccctgga
agccaccctgcgtctgtacctgcatccggaggcactgtccgaaaaactgccgactctgcgtctgctgac
tcgttccgcagaggttatccagatccaggcacagcgtctgcaagcaccgctggctgctcattacggtgc
ggagttcgcggtccaggtaatgccgtgtctgagccaaatcggttccggttctctgccggtagaccgtct
gccgagcgccgctctgactttcaccccgcacgatggtcgtggcagccacctggaatccctggcggcccg
ttggcgtgagctgccggttccggtaattggccgtatttacgatggtcgtctgtggctggacctgcgttg
cctggaagatgaacagcgctttctggagatgctgctgaaataatcctatcgccactttcagccaaaaaa
cttaagaccgccggtcttgtccactaccttgcagtaatgcggtggacaggatcggcggttttcttttct
cttctcaacacccttcgcgtcaacacttttccggctgccaaaccagatgtcaacacagctacaaaaaa
aggttagctcagtgcgaaagcactgagctaacctttttttaacgtccctatcagtgatagagattgac
atccctatcagtgatagagatactgagcacatcagcaggacgcactgaccggagagctgtcaccggatg
tgctttccggtctgatgagtccgtgaggacgaaacagcctctacaaataattttgtttaagggcccaag
ttcacttaaaaaggagatcaacaatgaaagcaattttcgtactgaaacatcttaatcatgcggattcta
gttaaatatgtcagaaaattcaatccgtctgactcaatattctcacggcgccggctgcggttgcaaaat
ttctccgaaagtgctggaaaccattctgcactccgaacaagcgaaatttgtagacccgaacctgctggt
aggtaacgaaacgcgcgacgacgccgcggtgtatgatctgggtaatggcacgtctgttatcagcactac
tgatttcttcatgccgatcgttgataacccgtttgatttcggccgtatcgctgcgaccaacgctatctc
cgatatctttgcaatgggtggtaagccgatcatggcgatcgcaattctgggttggccgatcaacaaact
gtccccggaaatcgcgcgtgaagttactgaaggtggtcgttatgcatgtcgtcaggcgggcattgcgct
ggcgggtggtcactctatcgatgcaccggagccaattttggtctggccgtgaccggtatcgtaccaac
cgaacgtgttaaaaagaactctaccgctcaggcgggttgcaaactgttcctgactaagccgctgggcat
cggcgttctgaccactgccgaaaaaaaatctctgctgaaaccggaacaccagggcctggccacggaagt
gatgtgccgtatgaacattgccggcgcttctttcgcaaatattgaaggtgtcaaagctatgaccgatgt
gactggcttcggcctgctgggtcacctgtctgagatgtgtcagggcgccggcgttcaggcccgtgtgga
ttatgaggcgatcccgaaactgccgggtgtggaggagtacattaaactgggtgcggttccgggcggcac
tgaacgtaacttcgcgagctacggtcatctgatgggtgaaatgcctcgtgaagtgcgcgatctgctgtg
tgatccgcagaccagcggcggtctgctgctggcggtgatgccggaagccgaaaacgaagttaaagcaac
cgccgcggaatttggtatcgaactgactgccatcggcgagctggtgccggctcgtggtggccgtgctat
ggtggaaatccgttaaatccggtttcatggagcctctggttcatctccggcaattaaaaaagcggctaa
ccacgccgctttttttacgtctgcaggacgtgctggatgcacacactggcttaagatgacaaaaaaaac
cattacacagtgcgaaagcactgtgtaatggtttttttaacgaaaatatatttttcaaaagtatcgtt
taccgctagctcagtcctaggtacaattacagccatcgtacgagcccggagggaagatggtcgtccccg
gtggggcggctggactctaaatccagttggggccgccagcggtcccggtcaggttcgactccttgcatc
ttccgccaatttgaaccccgcttcggcggggtttttttgttttctgtacatttcgtcaccctcccttcg
caataaacgcccgtaatagctggggacacacactggcttaagatgacaaaaaaaaggtgagttcagtgc
tttcgcactgaactcaccttttttttaacgtccctatcagtgatagagattgacatccctatcagtgat
agagatactgagcacatcagcaggacgcactgaccggagagctgtcaccggatgtgctttccggtctga
tgagtccgtgaggacgaaacagcctctacaaataattttgtttaagggcccaagttcacttaaaaagga
gatcaacaatgaaagcaattttcgtactgaaacatcttaatcatgctcaggccccttaaatatgcgact
gctgatactcaccggcccgcccgggtcaggtaaaacgtgtttcgcgcgggagctggctcggaactccg
gcaggaaggctggcgagtggcccatgtggaggcggacgctctccgcggattcctatgggatgagttcga
cccgaaactcgagcaagtggcccgggaactctttttgaagtccgttgaaacgtgtctggacgccgagct
agacctggtgatagcggacgacactaattactattccagcatgcggcgtgagttggctctcctggcctt
ggagaggaaagttccgtgggggatagtgtacctccgtaccggtttagatacgtgcctacgtaggaaccg
cgagcgaggtgaacctatcccggaggaggtcgttcgtagaatttatgataggtttgaaccacccgagcc
ggatcggtggtgggagcgtgcgaccttggttctagacgattctcgggtttccgaggaagttctcgagtt
cgtcgagtcgggtctacgcgtcgagaagcccaaaaagcgacgtcggcgcactgacccctcctccgtgaa
tgaagtcgacgtacgcacccgtcaggttatgggggagctgatgcgtcgactctccgagacaggcgcggc
tacccaagagctagggcgcaagctcagtgaactgcgccgtgagatagtgtcctccgtggaggatccgga
aaaggccgtccgggagttccggcgtcgggccgaggaggtgatccgggaatgcctgcatggagatgggtg

atccggtttcatggagcctctggttcatctccggcaattaaaaagcggctaaccacgccgctttttt
acgtctgcaggaacgtgctgagcagttacagagatgttacgaaccacgaaaaaaaaccaatttgcagtg
cgaaagcactgcaaattggttttttttgaattcgagctggatccagcttcctcagggccagtggcttc
tgtttctatcagctgtccctcctgttcagctactgacggggtggtgcgtaacggcaaaagcaccgccgg
acatcagcgctagcggagtgtatactggcttactatgttggcactgatgagggtgtcagtgaagtgctt
catgtggcaggagaaaaaaggctgcaccggtgcgtcagcagaatatgtgatacaggatatattccgctt
cctcgctcactgactcgctacgctcggtcgttcgactgcggcgagcggaaatggcttacgaacggggcg
gagatttcctggaagatgccaggaagatacttaacagggaagtgagagggccgcggcaaagccgttttt
ccataggctccgcccccctgacaagcatcacgaaatctgacgctcaaatcagtggtggcgaaacccgac
aggactataaagataccaggcgtttccccctggcggctccctcgtgcgctctcctgttcctgcctttcg
gtttaccggtgtcattccgctgttatggccgcgtttgtctcattccacgcctgacactcagttccgggt
aggcagttcgctccaagctggactgtatgcacgaaccccccgttcagtccgaccgctgcgccttatccg
gtaactatcgtcttgagtccaacccggaaagacatgcaaaagcaccactggcagcagccactggtaatt
gatttagaggagttagtcttgaagtcatgcgccggttaaggctaaactgaaaggacaagttttggtgac
tgcgctcctccaagccagttacctcggttcaaagagttggtagctcagagaaccttcgaaaaaccgccc
tgcaaggcggttttttcgttttcagagcaagagattacgcgcagaccaaaacgatctcaagaagatcat
cttattaatcagataaaatatttctagatttcagtgcaatttatctcttcaaatgtagcacctgaagtc
agccccatacgatataagttgtaattctcatgtttgacagcttatcatcgataagctttaatgcggtag
tttatcacagttaaattgctaacgcagtcaggcaccgtgtcggttgcgcctgagcaggaaaaaaaggt
ttagacagtgctttcgcactgtctaaaccttttttttatctatcgcgtacacagcgacagtcattcat
ctttctgcccctccaaaagcaaaaacccgccgaagcgggtttttacgtaaatcaggtgaaactgaccga
cgaactaaaacggatttaagacccactttcacatttaagttgtttttctaatccgcatatgatcaattc
aaggccgaataagaaggctggctctgcaccttggtgatcaaataattcgatagcttgtcgtaataatgg
cggcatactatcagtagtaggtgtttccctttcttctttagcgacttgatgctcttgatcttccaatac
gcaacctaaagtaaaatgccccactgcgctgagtgcatataatgcattctctagtgaaaaaccttgttg
gcataaaaaggctaattgattttcgagagtttcatactgtttttctgtaggccgtgtacctaaatgtac
ttttgctccatcgcgatgacttagtaaagcacatctaaaacttttagcgttattacgtaaaaaatcttg
ccagctttccccttctaaagggcaaaagtgagtatggtgcctatctaacatctcaatggctaaggcgtc
gagcaaagcccgcttattttttacatgccaatacaatgtaggctgctctacacctagcttctgggcgag
tttacgggttgttaaaccttcgattccgacctcattaagcagctctaatgcgctgttaatcactttact
tttatctaaacgagacatatttaaaacaccggcgcatgattaagatgtttcagtacgaaaattgctttc
attgttgatctccttttttaagtgaacttgggcccttaaacaaaattatttgtagaggctgtttcgtcct
cacggactcatcagaccggaaagcacatccggtgacagctctccgggctcgtacgatggctgtaattgt
acctaggactgagctagctgtaatcgaacttttcgccggtcgacagccactcaactccagcatgagatc
cccgcgctggaggatcatccagccggcgtcccggaaaacgattccgaagcccaacctttcatagaaggc
ggcggtggaatcgaaatctcgtgatggcaggttgggcgtcgcttggtcggtcatttcgaaccccagagt
cccgctcagccaatcgactggcgagcggcatcgcattcttcgcatcccgcctctggcggatgcaggaag
atcaacggatctcggcccagttgacccagggctgtcgccacaatgtcgcgggagcgaatcaaccgagca
aaggcatgaccgactggaccttccttctgaaggctcttctccttgagccacctgtccgccaaggcaaag
cgctcacagcagtggtcattctcgagataatcgacgcgtaccaacttgccatcctgaagaatggtgcag
tgtctcggcaccccatagggaacctttgccatcaactcggcaagatgcagcgtcgtgttggcatcgtgt
cccacgccgaggagaagtacctgcccatcgagttcatggacacgggcgaccgggcttgcaggcgagtga
ggtggcaggggcaatggatcagagatgatctgctctgcctgtggccccgctgccgcaaaggcaaatgga
tgggcgctgcgctttacatttggcaggcgccagaatgtgtcagagacaactccaaggtccggtgtaacg
ggcgacgtggcaggatcgaacggctcgtcgtccagacctgaccacgagggcatgacgagcgtccctccc
ggacccagcgcagcacgcagggcctcgatcagtccaagtggcccatcttcgaggggccggacgctacgg
aaggagctgtggaccagcagcacaccgccgggggtaaccccaaggttgagaagctgaccgatgagctcg
gcttttcgccattcgtattgcatatgtatatctccttcttaattaattaggatccgtgcccgctgaact
tcttcgtcacttactttagcaaaattagtgcccttcgggaaaaaatccctgaccaatccattcgtattc
tcattcgacccacgctgccacggcgaatgatgatcagaattccggctgttccgagcgaatcgctggacg
tgtacacctgtaccaaccactagttgattagctaagccgtatatatgagtaaacttggtctgacagaaa

gcaagctgataaaccgatacaattaaaggctcctttggagccttttttttttggagattttcaacatga
aaaaattattattgcggccgcccctgaattcgcatctagactgaactggccgataattgcagacgaaaa
aaaaggttgtatcagtgctttcgcactgatacaaccttttttttaacg
